# Pial surface CSF-contacting texture, subpial and funicular plexus in the thoracic spinal cord in monkey: NADPH diaphorase histological configuration

**DOI:** 10.1101/2020.01.30.927509

**Authors:** Yinhua Li, Wei Hou, Yunge Jia, Xiaoxin Wen, Chenxu Rao, Ximeng Xu, Zichun Wei, Lu Bai, Huibing Tan

**Affiliations:** Department of Anatomy, Jinzhou Medical University, Jinzhou, Liaoning 121001, China; Department of Neurobiology, Jinzhou Medical University, Jinzhou, Liaoning 121001, China

**Keywords:** NADPH diaphorase, Thoracolumber spinal cord, Non-human primate, funicular plexus, subpial plexus, cerebral spinal fluid

## Abstract

In spinal cord, white matter is distinguished from grey matter in that it contains ascending and descending axonal tracts. While grey matter gets concentrated with neuronal cell bodies. Notable cell bodies and sensory modality of cerebral spinal fluid (CSF) in white matter are still elusive in certain segment of the spinal cord. Monkey Spinal cord was examined by NADPH diaphorase (NADPH-d) histochemistry. We found that NADPH-d positive neurons clustered and featured flat plane in mediolateral funiculus in caudal thoracic and rostral lumber spinal cord, especially evident in the horizontal sections. Majority of NADPH-d funicular neurons were relatively large size and moderately-or lightly-stained neurons. In horizontal section, the multipolar processes of the neurons were thicker than that of regular other neurons. The processes oriented laterally or obliquely in the lateral funiculus. Some of neuronal cell bodies and proximal processes attached NADPH-d positive buttons or puncta. The neuronal processes interlaced network medially linked to lateral horn (intermediolateral nucleus, IML) and laterally to subpial region, in which formed subpial plexus with subpial NADPH-d neurons. Subpial plexus appeared to contacting externally with CSF. The subpial plexus patterned like round brackets located in lateromarginal pial surface. Compared with sympathetic IML in rostral thoracic segments and sacral parasympathetic IML, the funicular plexus configurated a specialized neuro-texture in caudal thoracic segments. The dendritic arbor of funicular neuron featured variety geometric plane shapes. The funicular plexus oriented exclusive layered flat-plane organization between lateral horn and subpial region in caudal thoracic and rostral lumber spinal cord. The subpial plexus may work as CSF sensor outside of spinal cord. The cluster of funicular neurons may function as locomotion sensor, besides visceral regulation. Different to periventricular CSF contacting or pericentral canal structures, NADPH-d funicular neurons and subpial plexus that located in the pial surface. With advantage of NADPH-d, we found funicular neurons which termed academically as funicular plexus and specialized localization for subpial structure we termed subpial plexus. The funicular texture was regarded as neuronal bridge between the interior CSF in the central canal and external CSF out of the pial surface.

## Introduction

Segmental arrangement is a typical organization for spinal cord[1]. The lateral horn[2] is usually referred to intermediolateral column (IML)[3] related to segmental variation for sympathetic component[4, 5]. Similarly, white matter and gray matter are two typical components in the spinal cord. White matter mainly consists of ascending and descending fibers. While, grey matter consists of neuronal cell bodies, axon terminals, and dendrites. Architecture of the neuronal cell bodies in the grey matter is characteristic of Rexed’s laminae[6–8]. Subpopulations of sympathetic neurons can be detected and also vary innervation of several neuropeptides in the thoracolumbar spinal cords [9, 10]. The nicotinamide adenine dinucleotide phosphate-diaphorase (NADPH-d) reaction is used as a marker to characterize certain neuronal properties and colocalize with nitric oxide synthase (NOS) [11, 12]. NADPH-d positive neurons have been detected in the spinal cord[13, 14].The distribution of NADPH-d neurons are majorly detected in superficial dorsal horn, deeper laminae dorsal horn, Laminae X and IML. NADPH-d neurons is scarcely found in ventral horn in rat, but can be detected in the dog[15] and pigeon[16]. Occasionally, positive neurons are detected in the white matter[14]. However, the occurrence of neuronal cell bodies is individually detected in white matter in some species, such as pigeon[16, 17]. Torresdasilva demonstrates that the distribution of NADPH-d neurons in non-human primate is similar to other species[18]. In development study of human, NADPH-d neurons and fibers are well-defined the sympathetic and parasympathetic neurons in the IML and the lateral collateral pathway and the medial collateral pathway (MCP)[19]. Dun et al reports that NOS-positive fibers are detected in the thoracic white matter of rat, mouse, cat and squirrel monkey[20]. The distribution of NOS-positive fibers is more than that of other segments. However, distribution of NADPH-d neurons and fibers is still lack in thoracolumbar spinal cord of non-human primate species.

Referred to regional cytoarchitecture in developing of cortex[21], cat spinal cord[22] and a previous investigation of funiculus neurons in thoracic spinal cord in non-human primate species[23], the pattern of neuronal distribution in our present data could be partitioned into several anatomical and functional areas: subependymal zone, nucleus intercalatus spinalis, intermediolateral nucleus(column), funicular zone, subpial zone. The similar results are partially verified in human fetal spinal cord[24]. Autonomic innervation of four subnuclei in thoracic spinal cord to intermediolateralis pars funiculus (IMPf) in cat [22], rat[22]. There is funicular neuron found in earlier investigation of cyto-and fiber-architecture of the intermediolateral nucleus of cat spinal cord[25].

Recently, we also investigated NADPH-d positivity in non-human primate. We found a specialized NADPH-d neuronal organization in the lateral funiculus in caudal thoracic and rostral lumbar spinal cord. To the best features of NADPH-d staining for the neuroanatomy, it reveals high quality of dendritic arborization of neurons. The cluster of NADPH-d neurons formed ganglion-like neuronal arrangement lateral IML with their processes appeared to reach subpial region and contacting cerebral spinal fluid (CSF). Usually, CSF contacting neurons are neurosecretory cells with an apical extension into the central canal[26, 27]. It is GABAergic PKD2L1-expressing neurons detected in zebrafish, mouse and macaque[28]. The present mentioned CSF contacting neurons located in the white matter connected in the grey matter. Our new finding of CSF contacting neuron may send several processes to spinal surface. It also looked like mechanical device. The aim of the investigation is to demonstrate CSF contacting circuit in lateral funicular location of the thoracolumbar spinal cord in non-human primate.

## Materials and Methods

Twelve monkeys were used for the present investigation. All animal care and treatment in this study were performed in commercialized animal facilities for scientific research accordance with the guidelines for the national care and use of animals approved by the National Animal Research Authority (P.R. China). The ambient temperature was maintained at 24±2 °C and relative humidity at 65±4%. Reverse osmosis water was available *ad libitum*. Normal food, fresh fruit and vegetables were supplied twice daily. Animals were purchased from Beijing Xierxin Biological Resources Research Institute (Beijing, China), Hainan Jingang Biotech Co., Ltd (Hainan, China) and Wincon TheraCells Biotechnologies Co., Ltd. (Naning, China). All experimental procedures were approved by the Institutional Ethics Committee in Animal Care and Use of the Jinzhou Medical University.

All the tissues were obtained in the lab of the experimental monkey companies from Xierxin Biological Resources Research Institute and Wincon TheraCells Biotechnologies Co. Ltd, respectively. Animals were deeply anaesthetized by the professional technicians with pentobarbital sodium and then intracardiacally perfused with normal saline followed by paraformaldehyde 4% in 0.1 M phosphate buffered saline (PBS, pH 7.4). After perfusion, spinal cords cervical to coccygeal and were removed and postfixed in 4% paraformaldehyde overnight at 4 °C. The tissues were next transferred into 25% sucrose in 4% paraformaldehyde PBS for cryoprotection. Spinal cord segments were sectioned either coronally or horizontally at 40 μm using a cryostat (Leica).

### NADPH-d histochemistry

The spinal cord sections were processed for N-d activity according to the procedure of previous study[29]. Briefly, sections were incubated in 0.1 M PBS pH 7.4, containing 0.3% Triton X-100, 0.5 mg/ml nitroblue tetrazolium (NBT, Sigma, Shanghai, China) and 1.0 mg/ml β-NADPH (Sigma, Shanghai, China) at 37°C for 1-3 h. Following the reaction, the sections were rinsed in PBS, air-dried and coverslipped with Neutral Balsa (China).

Images were captured with a DP80 camera in an Olympus BX53 microscope (Olympus, Japan). Sections were observed under the light microscope and randomly selected sections from all spinal cord levels in each animal were quantitated using Olympus image analysis software (Cellsens Standard, Olympus). All data are expressed as the mean ± SEM and P < 0.05 were regarded as statistically significant. Differences between funicular and IML neurons were analyzed using t-tests.

## Results

The neurons and neuronal texture were detected in white matter of the spinal cord by NADPH-d histological staining. The NADPH-d positive neurons generally distributed as in other species in the dorsal horn, intermediolateral horn (related termed IML column or nucleus), area around the central canal, dorsal commissural nucleus (DCN) and individually in the anterior(ventral) horn. Differently, NADPH-d positive neurons were also notably found in the white matter of the thoracolumbar segment. And tremendous neuronal processes were visualized appearance of fine texture. The size and shape of the cells and fibers, and intensity of staining varied in location. The cytoarchitecture for the IML and lateral funiculus in the white matter were focused in the present experiment and summarized the data by coronal section and horizontal sections.

We started to demonstrate the results in the transverse sections following with the horizontal sections. Many sections were so large field that we montaged several sections in following figures. Figure 1 and Figure 2 showed NADPH-d positivity in the transverse sections in the thoracolumbar segment. Besides regular neuronal and fibers in the gray matter, such as lateral horn, distinct neuronal texture was detected in the lateral funiculus. The Clarke’s nucleus was constitutively detected in the segments. Intensely NADPH-d-positive neurons were detected in the lateral horn. There were several distinct bulges showed in the neuronal texture in the lateral funiculus. Some neurons in the bulges were surround by fiber tract in the lateral funiculus. NADPH-d reactive fibers were revealed in the mediolateral white matter. The positive cells broadly distributed from the lateral horn to the pial surface. In the lateral horn, three sub-regions revealed: the dorsal subregion received input fibers from the lateral co-lateral pathway and output fibers to the dorsal lateral neurons; middle region of fiber tracts intermingled with the white matter and the ventral lateral region sent output fibers to the lateral funiculus (Figure 1-2 B and C). Transverse section showed the funicular plexus, fiber tract(bundle) and subpial terminals as well as subpial neurons, which formed subpial plexus. The cytoarchitecture showed as a pattern of a part of wheel spokes in the intermediate zoom of the lateral funiculus. The subregional classification helped to locat the funicular neurons and funicular plexus. The most of funicular textures occurred ventral to the IML and the dorsal subregion. Occasionally, spheroid was detected (Figure 1-2 I). One more example of the transvers section was figure 2-1 and figure 2-2. In the mid-lower lumber and sacral spinal cord, the typical funicular neurons were scarcely detected (Figure 3).

**Figure 1-1.**
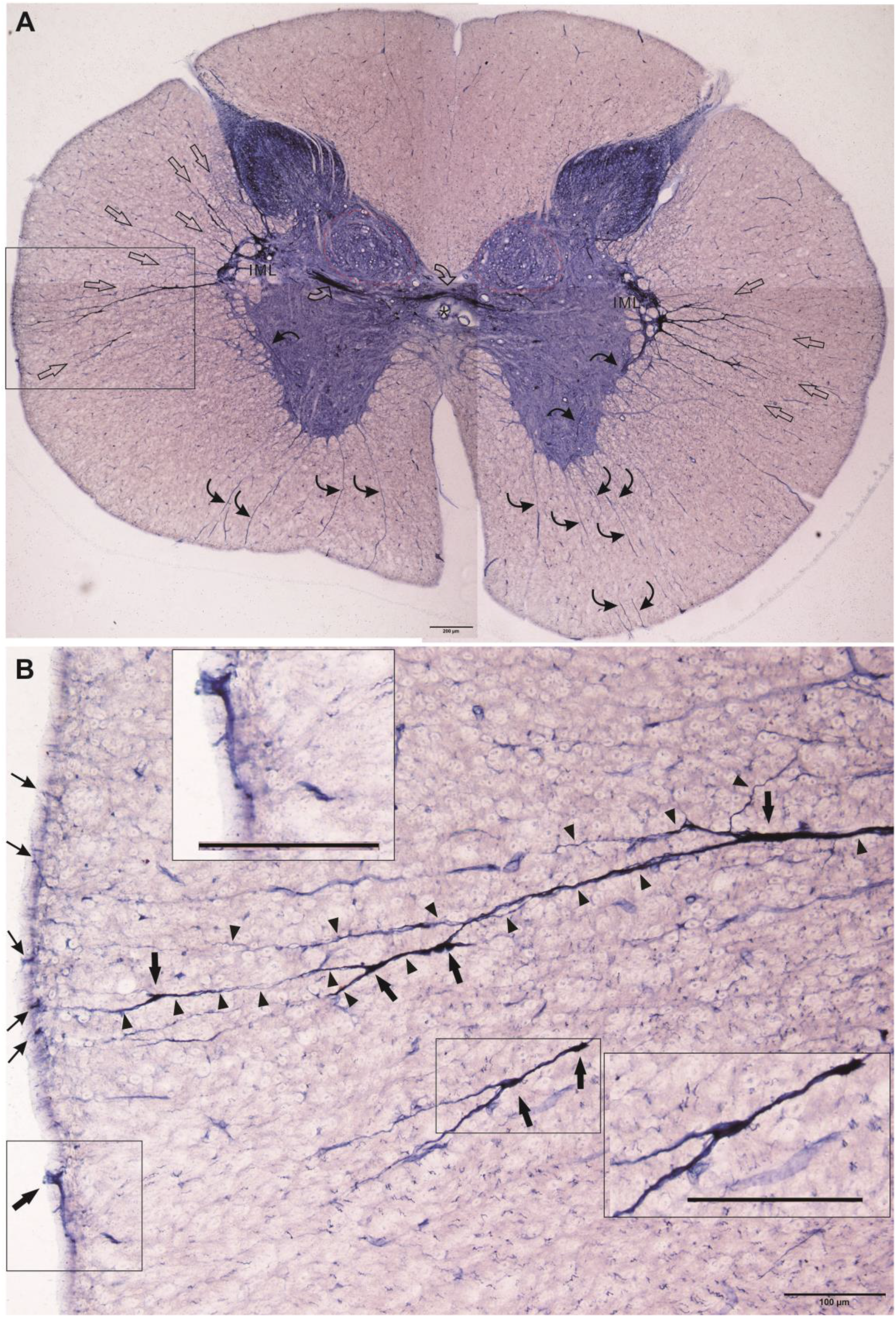
Subpial NADPH-d fiber and neuronal texture in the lateral funiculus in the thoracolumbar. A: Montage of sections showed the neuronal NADPH-d positivity. Two different patterns of NADPH-d positivity revealed in the IML of the lateral horn: impacted neuronal region and network fibers (plexus). Red dash line indicated the Clake’s nucleus. Open arrow indicated the radiated fibers directed to the pial surface. Curved arrow indicated NADPH-d tracts and fibers from IML to the central canal. Curved thin arrow indicated motor fibers traveling to anterior root. Asterisk indicated the central canal. B: Magnification from rectangle in A showed neuronal tracts and fibers in the mediolateral white matter. Arrowhead indicated one intact neuronal tract. Arrow indicated bulge on the tract in which may contain a neuron soma. Thin arrow indicated the puncta in the pial surface. Two insets showed neuronal tract or fiber and neuronal cells. Bar in A = 200μm and bar in B = 100μm.

**Figure 1-2.**
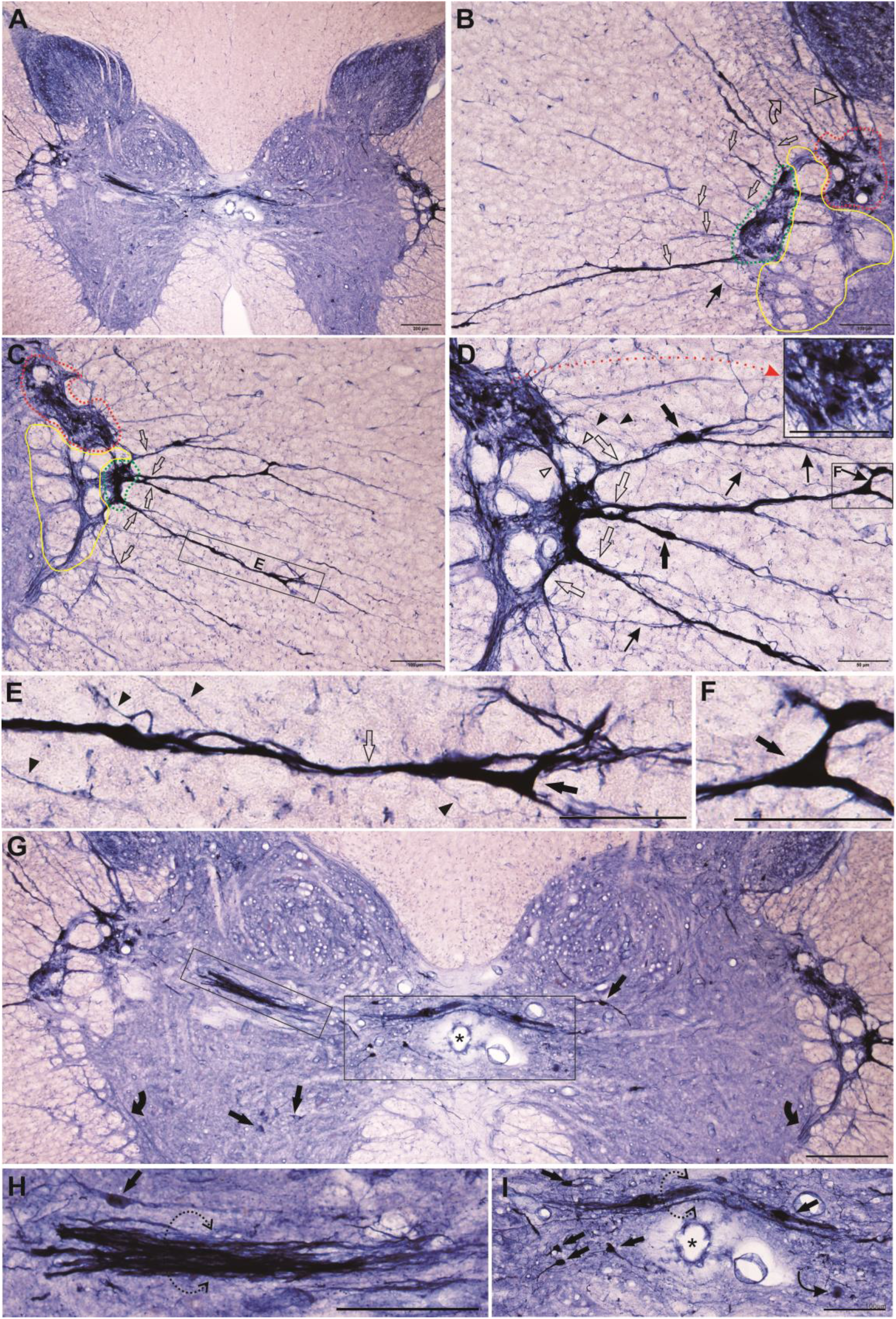
The detailed magnification of Figure 1-1A revealed the configuration of the lateral horn and IML in the gray matter and funicular texture in the white matter. A as guiding atlas for the following image showed similar the lateral horn and IML. B: Discernible three sub-regions showed in the magnification of the left side of the lateral horn. The dorsal subregion (red dash line) indicated the input fibers from the lateral co-lateral pathway (open arrowhead) and output fibers to the dorsal lateral neurons (curved open arrow). The sub-region of the yellow line indicated fiber tracts intermingled with the white matter. The ventral lateral region (green dash line) sent output fibers to the lateral funiculus. The open arrow indicated radiated bundles or tracts. Thin arrow indicated network fiber. C: the right-side lateral horn and Similar pattern to B. D: magnification from C. Inset showed some degenerated terminals (red dash arrow). The dorsal sub-region connected with the ventral lateral sub-region by network tracks (open arrowhead) and fibers(arrowhead). The fiber tracts (open arrow) also formed plexus by the fibers (thin arrow). The tracts contained neurons(arrow). E: Magnification from C. The bundle contained neuron (arrow) in along the thick fiber (open arrow) as well and the thin branching (arrowhead). F: magnification from D. Arrow indicated neuron. G: Thick track between the central canal and the lateral horn, and the dorsal of the central canal. Scattered neurons around the central canal and the anterior horn. H and I: magnification from G. Broken dash circle indicated fiber track. Arrow indicated neuron. Curved thin arrow indicated a spheroid in I. Bar in A = 200μm, bar in B, C and G-I= 100μm, bar in D-F = 50 μm

**Figure 2-1.**
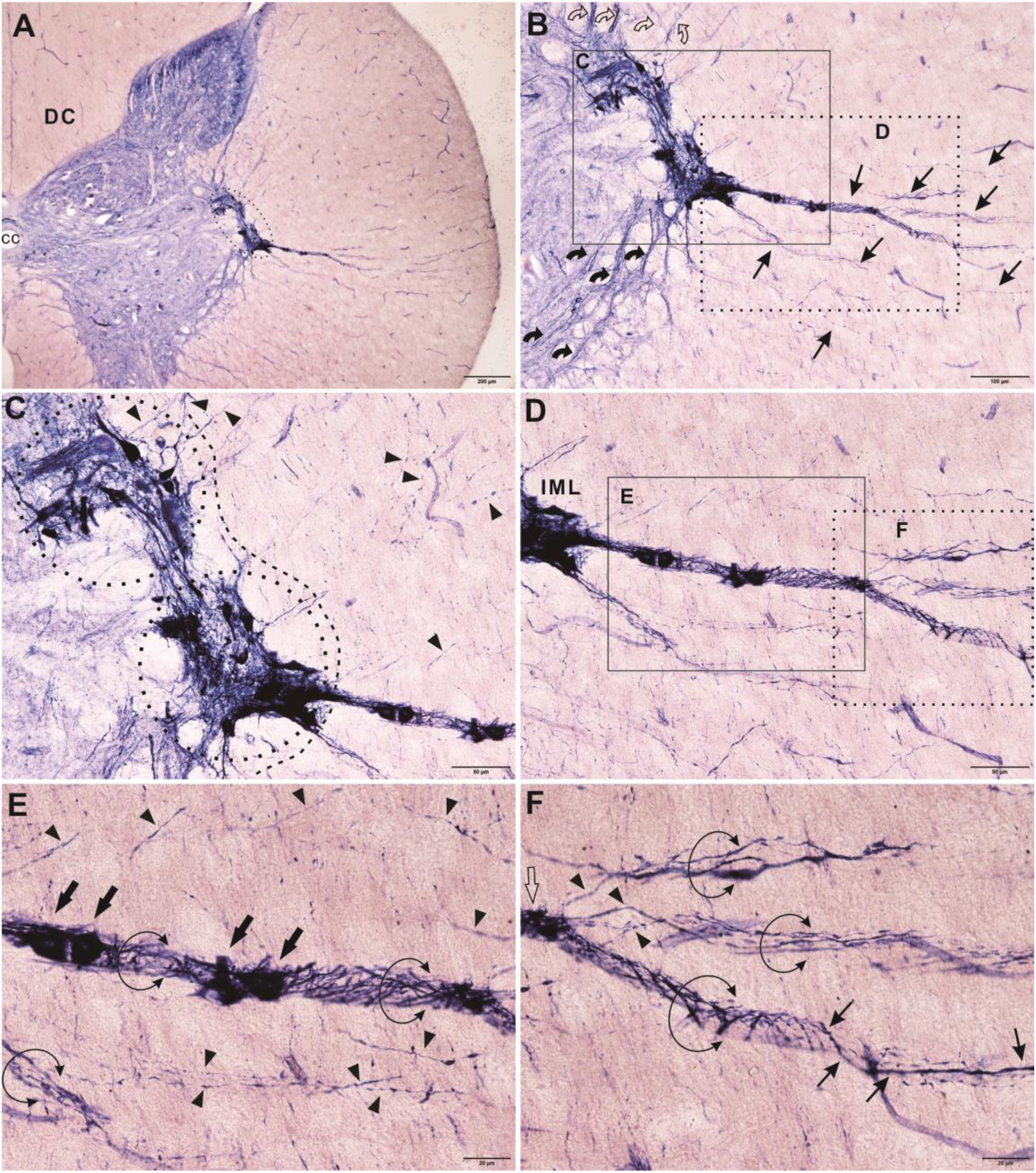
Transverse section showed the funicular plexus, fiber tract(bundle) and subpial terminals. A: Semi lateral of the Transverse section showed the gray matter and white matter. Dashed line indicated IML. DC for the dorsal column and cc for the central canal. B: The IML of the lateral horn and neuronal textures in the white matter. Curved arrow, curved open arrow and thin arrow indicated NADPH-d fibers of three directions. The rectangle C and D magnified following images. Curved arrow indicated the projecting fibers and bundles from the lateral horn. C: Dashed circle indicated two groups of subnuclei showed in the IML of the lateral horn. Dash line indicate the gray matter and white matter. Arrowhead indicated the fibers. D: The mediolateral funiculus and the neuronal texture from the IML. E: Magnification from D. Arrow indicated funicular neuronal soma. Open circle indicated funicular fiber plexus or fiber bundle. Arrowhead indicated single fiber. F: Magnification from dash rectangle in D. Three funicular plexus or fiber bundles. Arrowhead indicated the branch from one fiber plexus (open arrow). Bar in A = 200μm, bar in B = 100μm, bar in C and D = 50μm and bar in E and F = 20μm.

**Figure 2-2.**
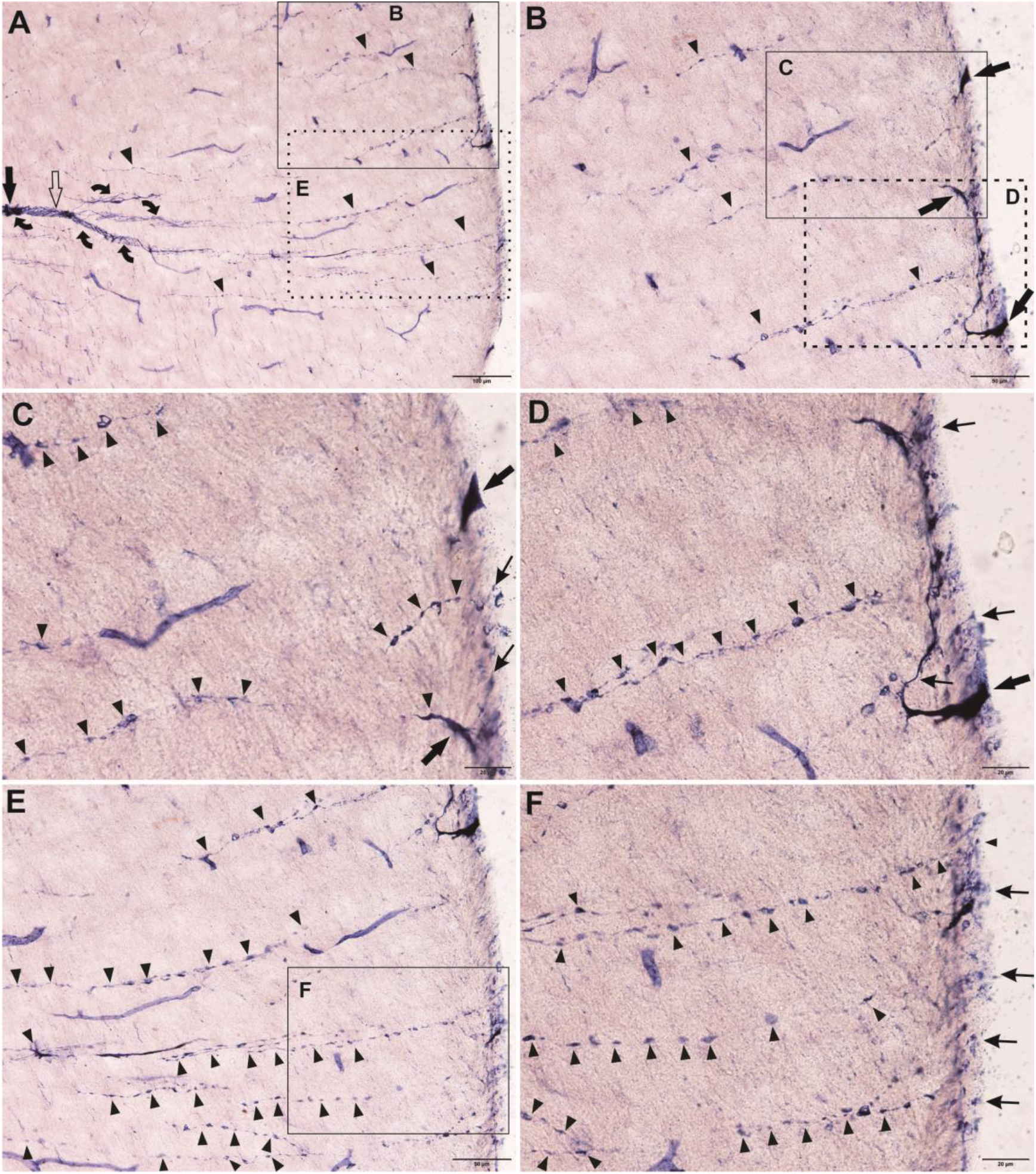
Transverse section showed the funicular fiber cord and subpial terminals (continued, lateral white matter to pial surface). A: Current figure was depicted on Figure 2-1F which both figure arrangement was created from the same section (Figure 2-1). Arrow indicated the funicular neuron in the fiber plexus (cord). Open arrow indicated the same region and the curved arrows indicated also three fiber cords in the last figure. The sector distribution of the fibers. Arrow head indicated the single fiber. Some of them reached to the pial surface. B: Magnification from A showed subpial fibers (arrowhead) and three subpial neurons (arrow) in the subpial region. C and D showed magnification from B. Thin arrow indicated the fibers in pial surface. Varicosity revealed in some fibers(arrowhead). E: Magnification from A showed the subpial fibers(arrowhead). F: Magnification from E showed some fibers reached to the pial surface (thin arrow). Bar in A = 100μm, bar in B and E = 50μm, bar in C, D and F = 20μm.

**Figure 3.**
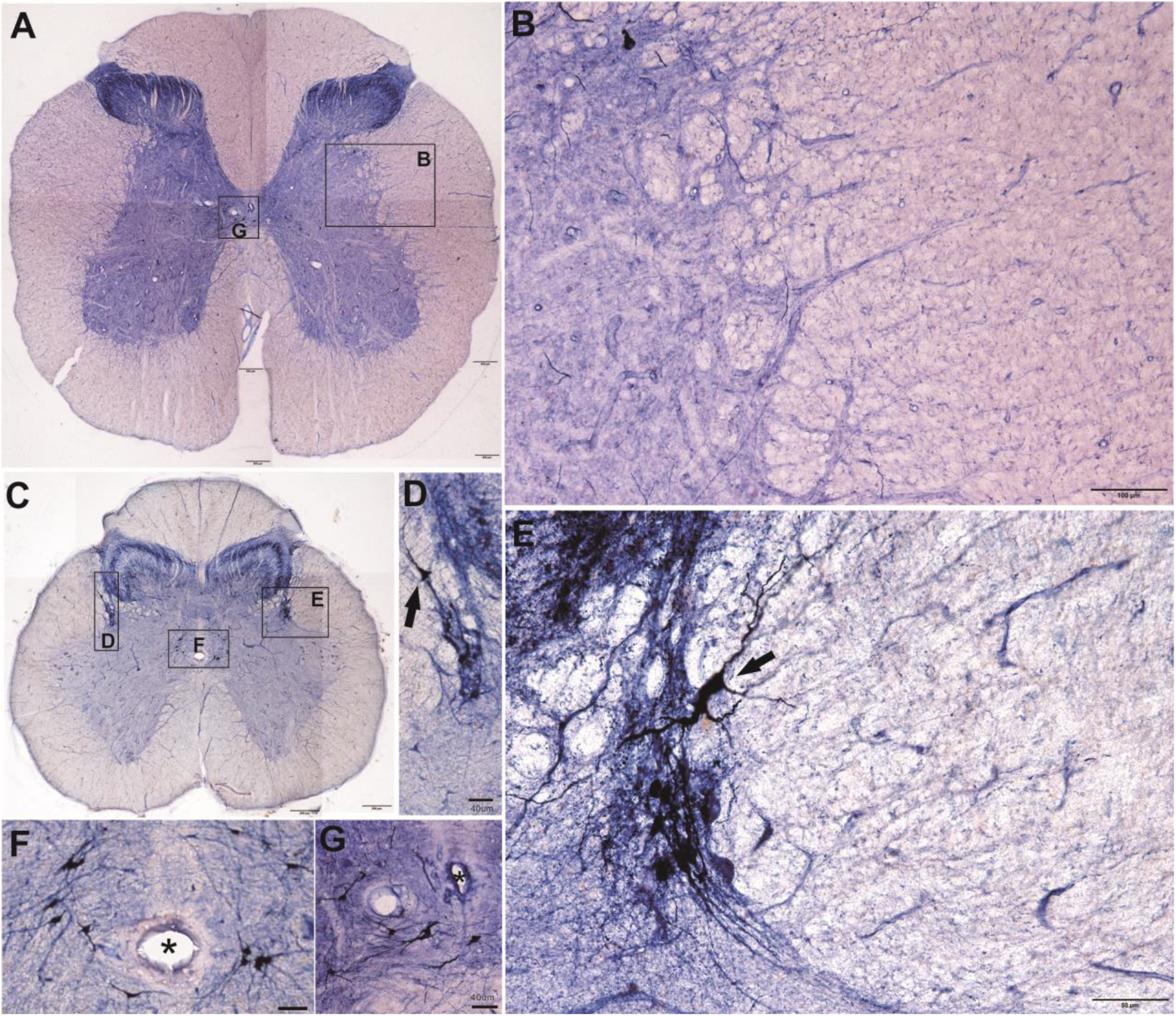
Transverse sections of the mid-lower lumbar and sacral spinal cord. Neurons were scarcely found in the lateral funiculus in the mid-lower lumber and sacral spinal cord A, B, C, D and E. Arrow indicated a neuron in the dorsal lateral nucleus (D) and the dorsal horn (E). F and G: magnification from A and C respectively showed the positive neurons around the central canal (asterisk). The ependymal cells showed NADPH-d positive reaction in G. Bar in A and C = 200μm, B= 100μm, D, E and G =40μm and E=50μm.

Horizontal sections showed flat plane neuronal plexus perpendicular to pial surface. NADPH-d staining showed neuronal texture in which the positive fiber cord intermingled with neuronal somas. We termed the neuronal texture as funicular plexus neurons funicular plexus. We also termed the neuronal texture in the subpial region and pial surface as subpial plexus. Firstly, we present a montage figure including both sides of pial surfaces and subpial structures through the central canal or central gray matter at the thoracic horizontal level (Figure 4). In the figure 4, the typical IML boundary with the solid line revealed the left half of the gray matter in upper side dorsal horn. At least three patterns of neuronal distribution were scattered in the lateral funiculus: individual cell, clustered neurons and ganglia-like cluster of neurons (Figure 5-1). According transverse section, the most of the typical funicular neurons and plexus located at the ventral level of the IML and dorsal subregion of the dorsal horn. So, we demonstrated the funicular neurons from the lower side of the white matter in Figure 4. Next, the Figure 5, 6 and 7 were all magnification from Figure 4. All of the funicular neurons majorly projected fibers to the IML in the lateral horn and pial surface beside pial neurons. The subpial neurons projected to the medially. Some dendrites of the funicular neurons were occasionally found running longitudinally (Figure 5-2).

**Figure 4.**
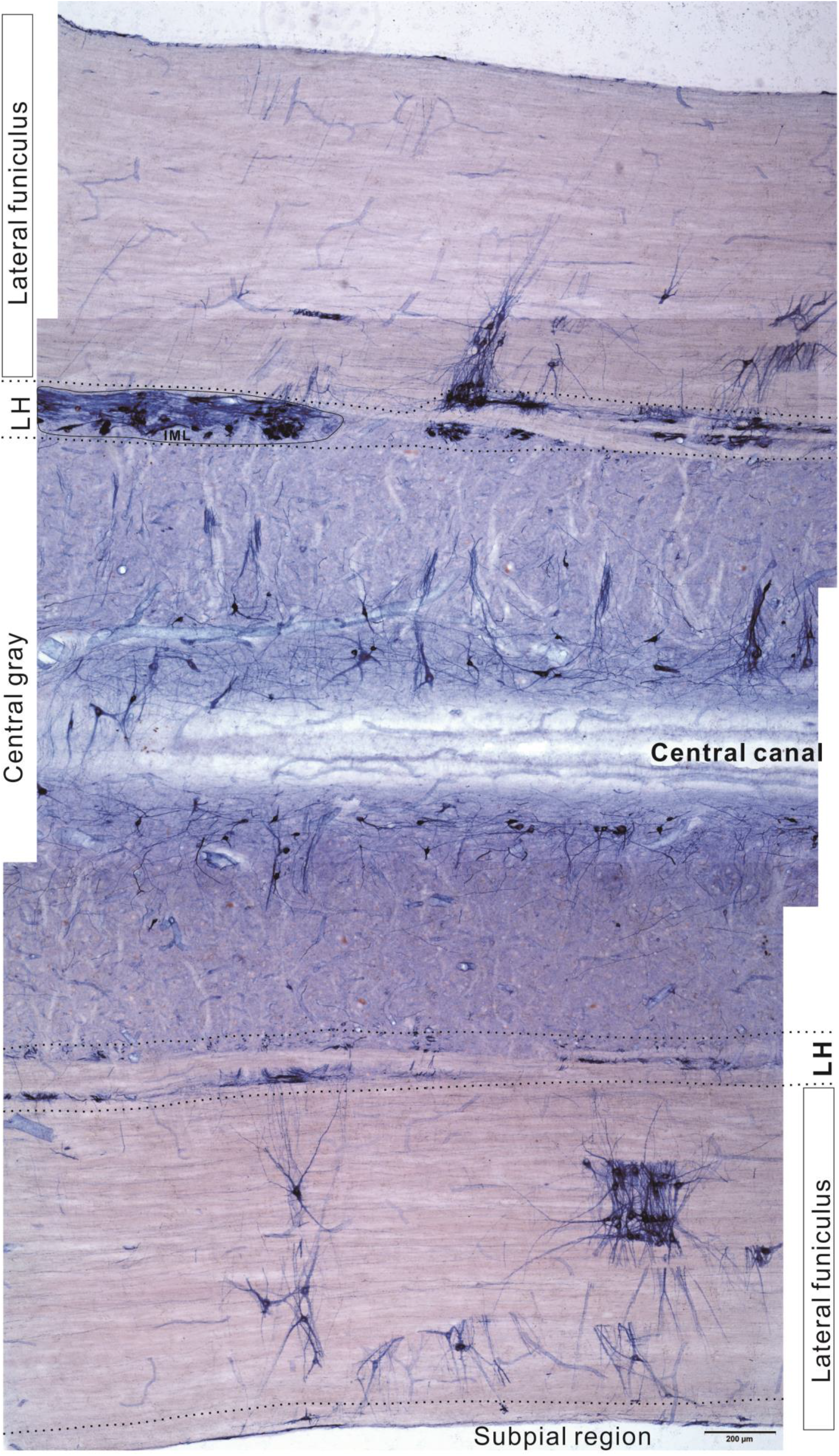
Montage of horizontal sections showed the subpial plexus, funicular plexus, IML (solid line), intercalated nucleus and central gray matter. Bar= 200μm

**Figure 5-1.**
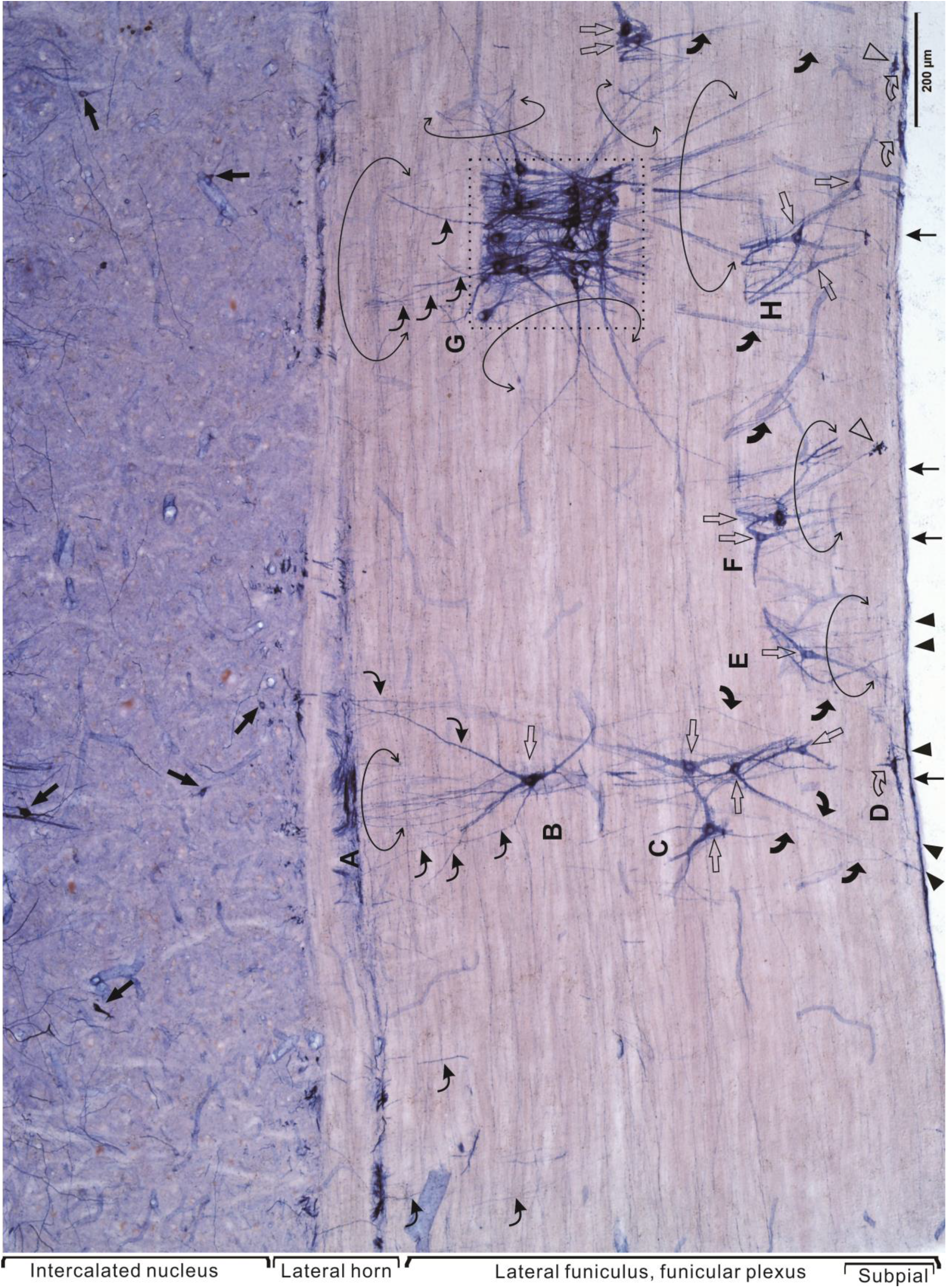
Enlarged view of the one side of lateral funiculus in the horizontal section from the previous image. Arrow indicated neuron in the gray matter, open arrow for the neuron in the white matter and curved open arrow for the neuron in the subpial region. Dash box revealed a ganglia-like plexus. Open circle indicated grouped fibers. Curved thin arrow indicated fiber projected to the lateral horn. Curved arrow indicated fiber injected to the subpial region. Arrowhead indicated fiber reached to the pial surface. Thin arrow indicated the fiber along pial. Open arrowhead swelling terminal. Bar= 200μm

**Figure 5-2.**
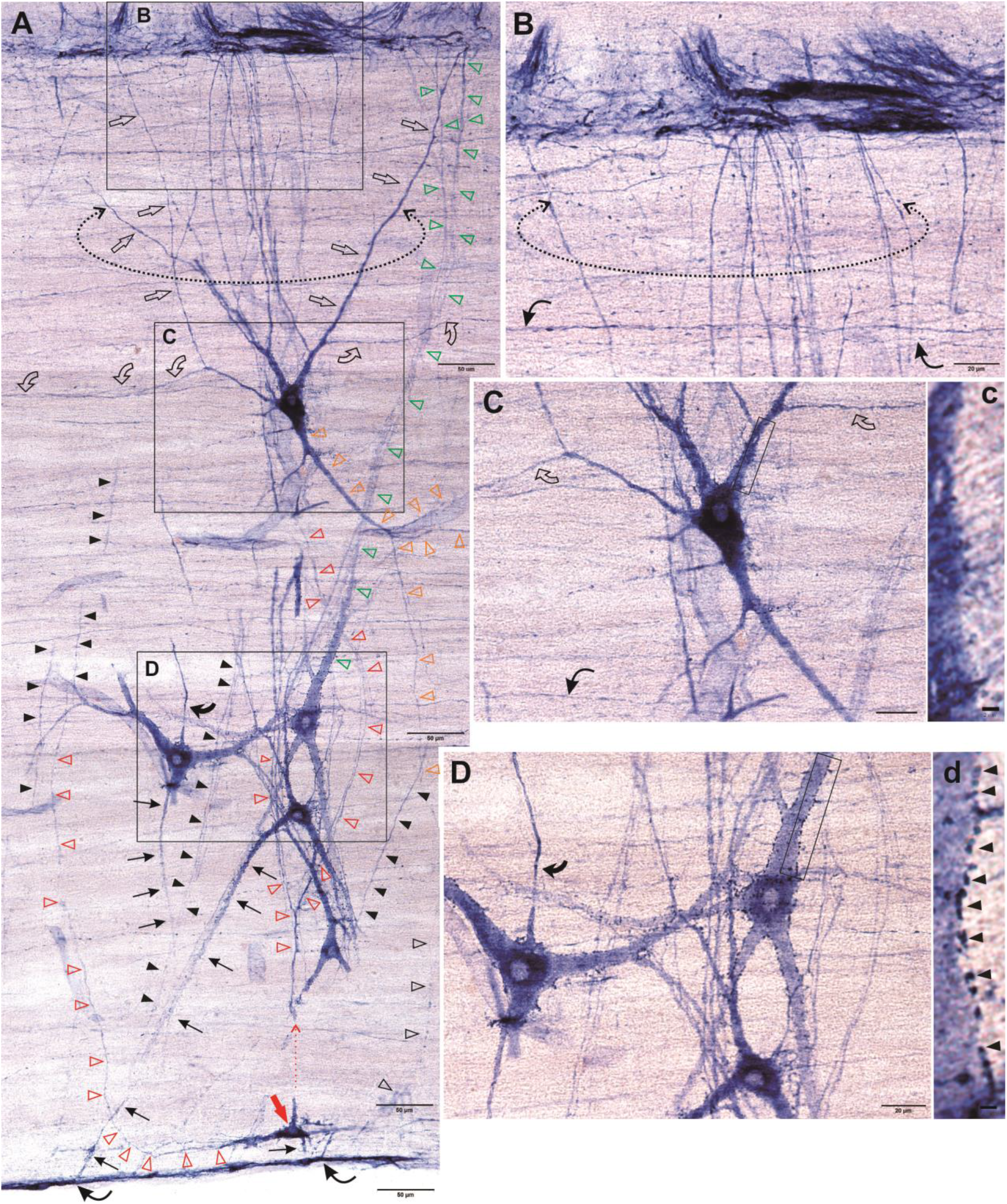
The funicular neurons and fibers as well as membrane particles surround neurons in the lateral funiculus. A: magnification from the location A, B, C and D in the previous image. Image A showed the divergent dendritic arbor and projected to the lateral horn (dash circle and open arrow). Arrow indicated a subpial neuron. Colored open arrows indicated fibers, red and green one to the gray matter, and green one to the pial surface. Arrowhead or black open arrowhead indicated the perpendicular fibers. Curved open arrow indicated longitudinal fibers of the funicular neurons and the same dendrites in C. Curved arrow indicated apical axon. Thin arrow indicated the fibers of projecting to the pial surface. Curved thin arrow indicated fibers reaching to the pail surface fiber. In B and C, curved thin arrow indicated longitudinal fibers. Compared C and D as well as both magnifications of c and d, the neurons of pericellular particles revealed in D. Bar in A = 50 μm, bar in B, C and D = 20μm and bar in c and d = 2μm.

Clustered punctate (arrows) were peri-cellularly associated with the large size multiple dendritic and lightly stained neurons (Figure 5-2, 5-3, 5-4 and 5-5 as well as Figure 7-2). The neuron of pericellular particles scattered individually or clustered. The pericellular particles could located membrane of soma and thick proximal dendrites. It also aggregated around soma (Figure 5-5). It may be aggregated NADPH-d molecules. The typical IML was detected in the Figure 6. As previous reports, the NADPH-d neurons and fibers also distributed in the IML, intercalated nucleus and central gray matter (Figure 6). It showed a relative morphological diversity of these neurons. There were large size multi-dendritic neurons in the intercalated nucleus which was similar funicular neurons (Figure 6B). The funicular neurons and plexus as well as subpial region texture also confirmed in the other side white matter of the same horizontal section (Figure 7-1 and 7-2). Together with above figures, the funicular neurons in funicular plexus was large size neurons compared with the IML neurons. The most of IML neurons were ovary shaped neuron, while the funicular neurons in funicular plexus were multiple dendritic neuron. The branching of the funicular plexus reduced the fiber cord size from the medial portion (proximal to the IML in the lateral horn) to the lateral terminals in the subpial region. The sector distribution of the fibers divergently spread a broad field toward the pial surface. The subpial plexus located along the surface of the subpial region. The subpial neuronal somas were aligning under pial surface (Figure 7-1 C and D).

**Figure 5-3.**
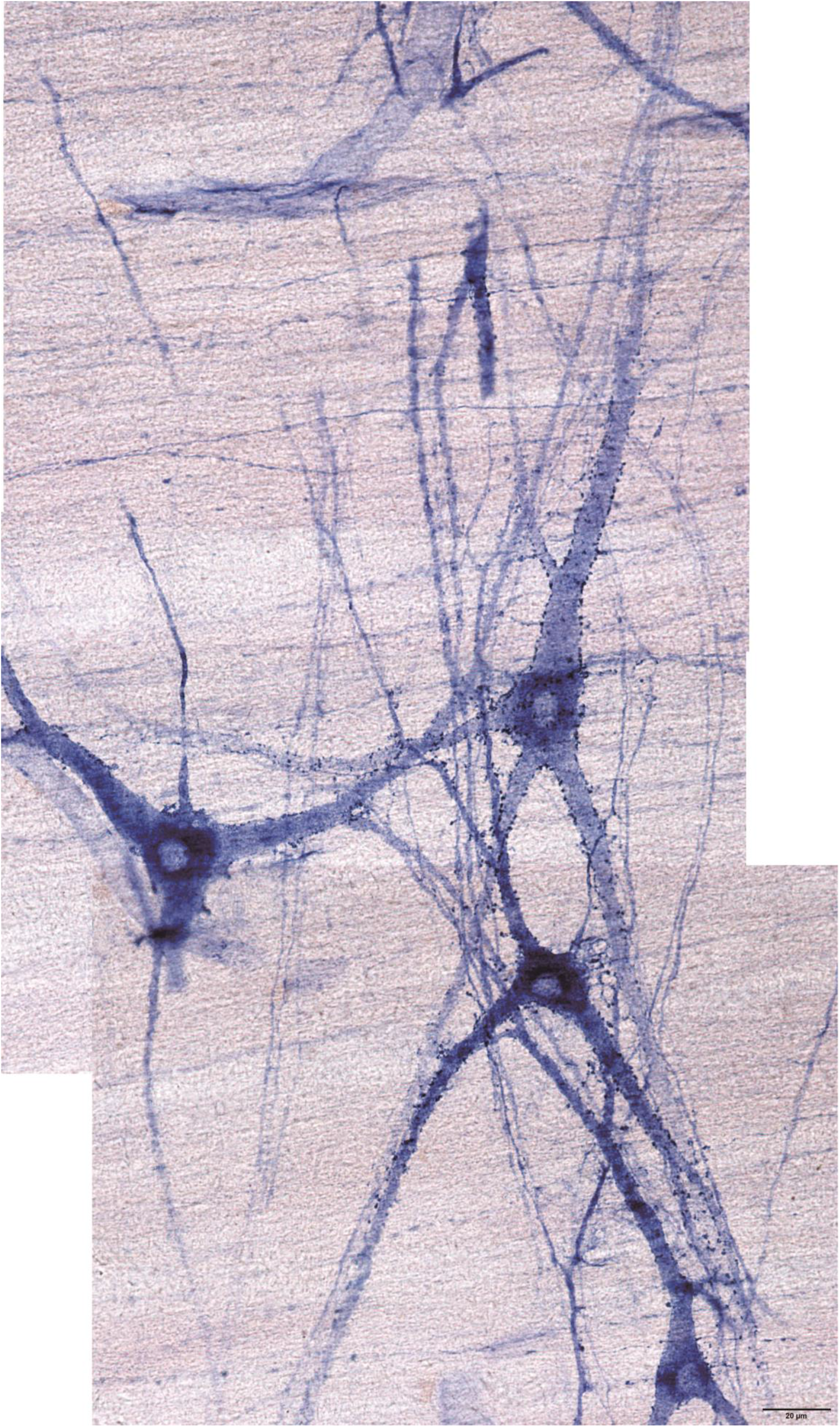
It was montage image. High power view of the previous image showed the neurons of the pericellular particles. Bar =20μm.

**Figure 5-4.**
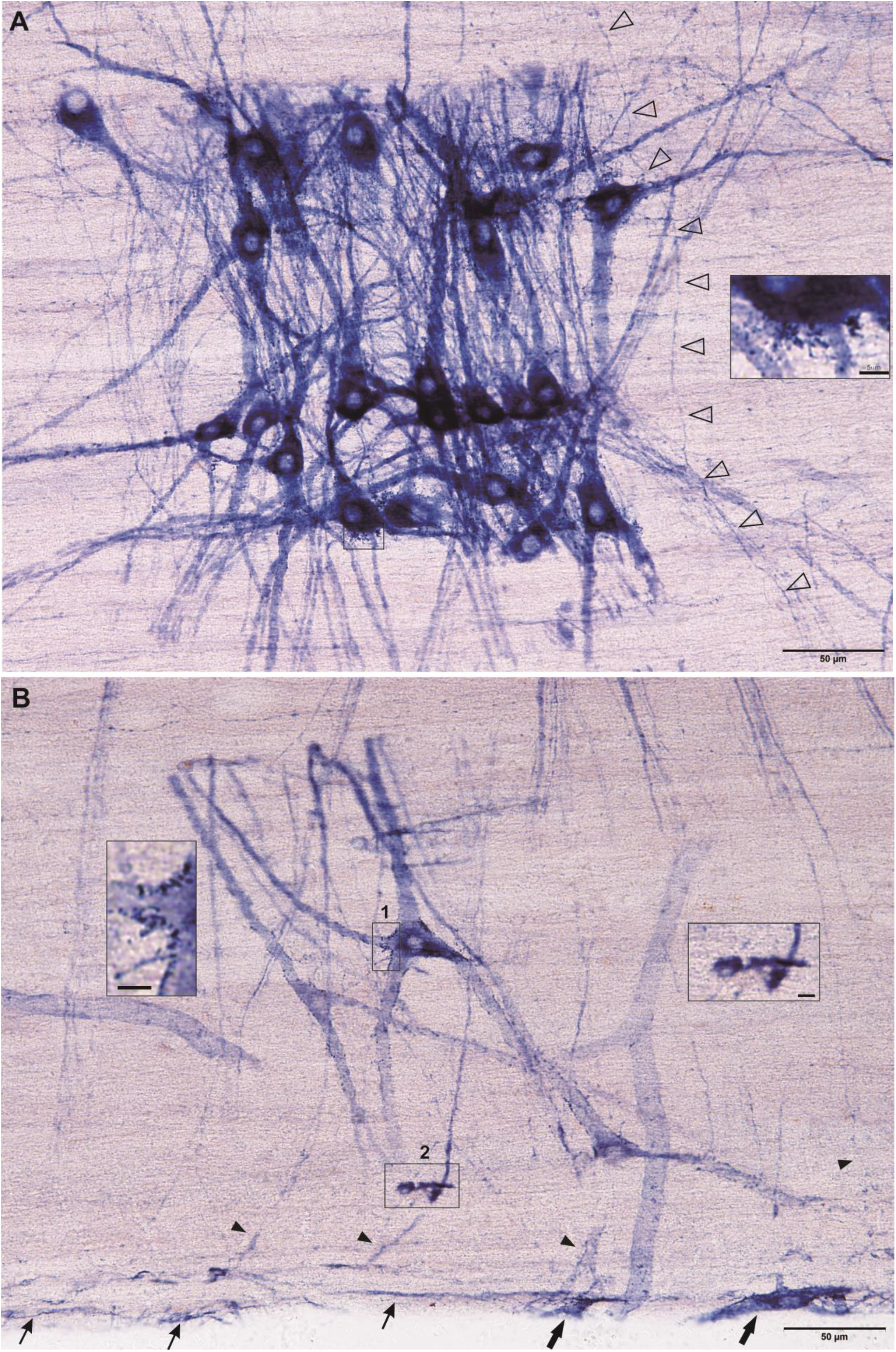
High power view from the previous image of the funicular neurons and fibers. A: Ganglia-like neurons. Inset showed pericellular particles (A and square 1 in B). Open arrowhead indicated passing fiber. B: Arrow indicated neurons located in the pial surface. Thin arrow indicated fibers located in the pial surface. Irregular degeneration from in square 2. Bar A and B = 50μm. Bar for inset = 5μm

**Figure 5-5.**
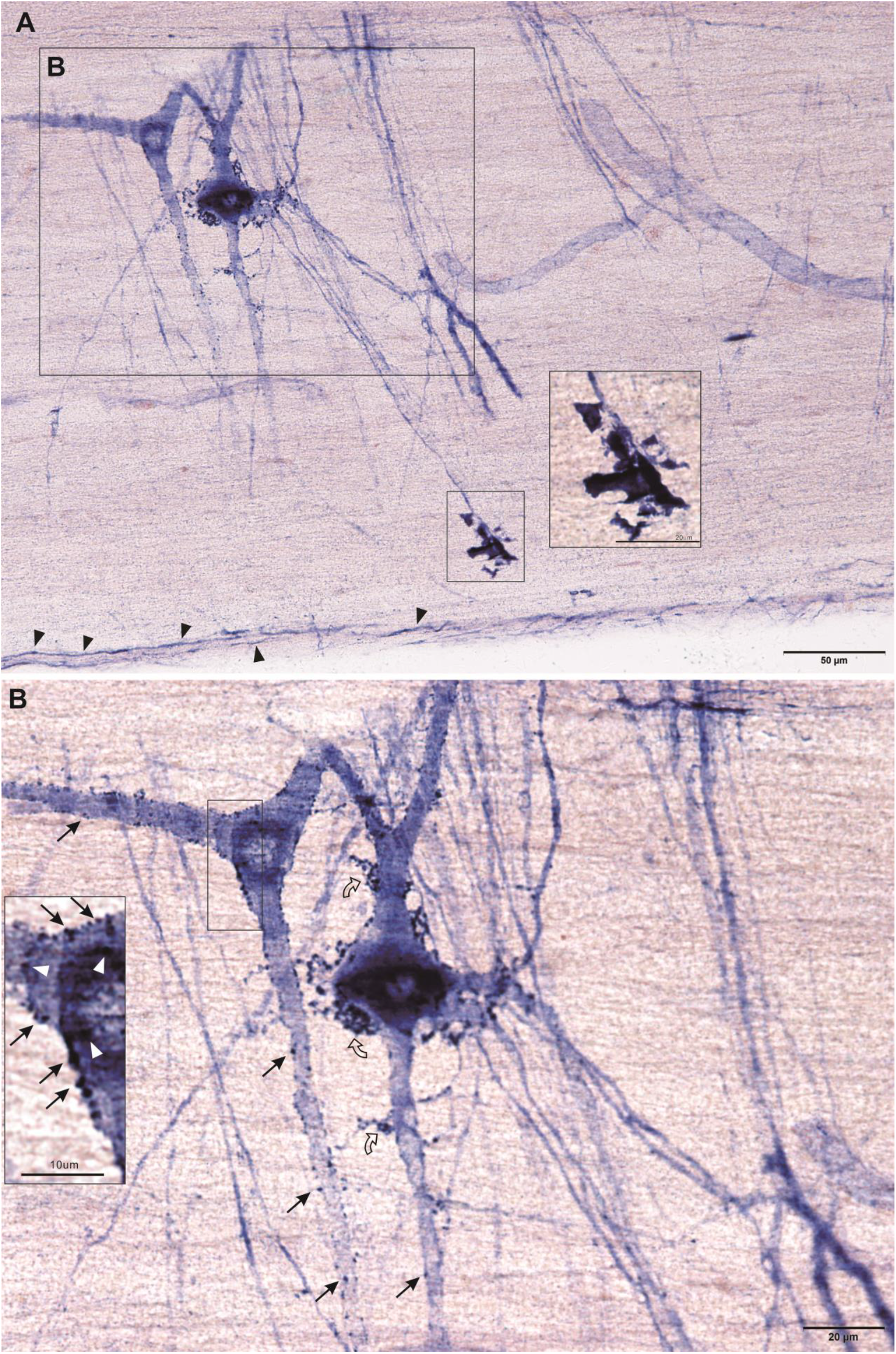
High power view from the previous image of the funicular neurons and fibers. A: subpial fiber(arrowhead) and irregular degeneration (inset). B: Magnification of B. Thin arrow indicated pericellular particles. Curved open arrow indicated clustered particles. Inset showed intracellular particles (white arrowhead). Bar in A= 50μm, in B

**Figure 6.**
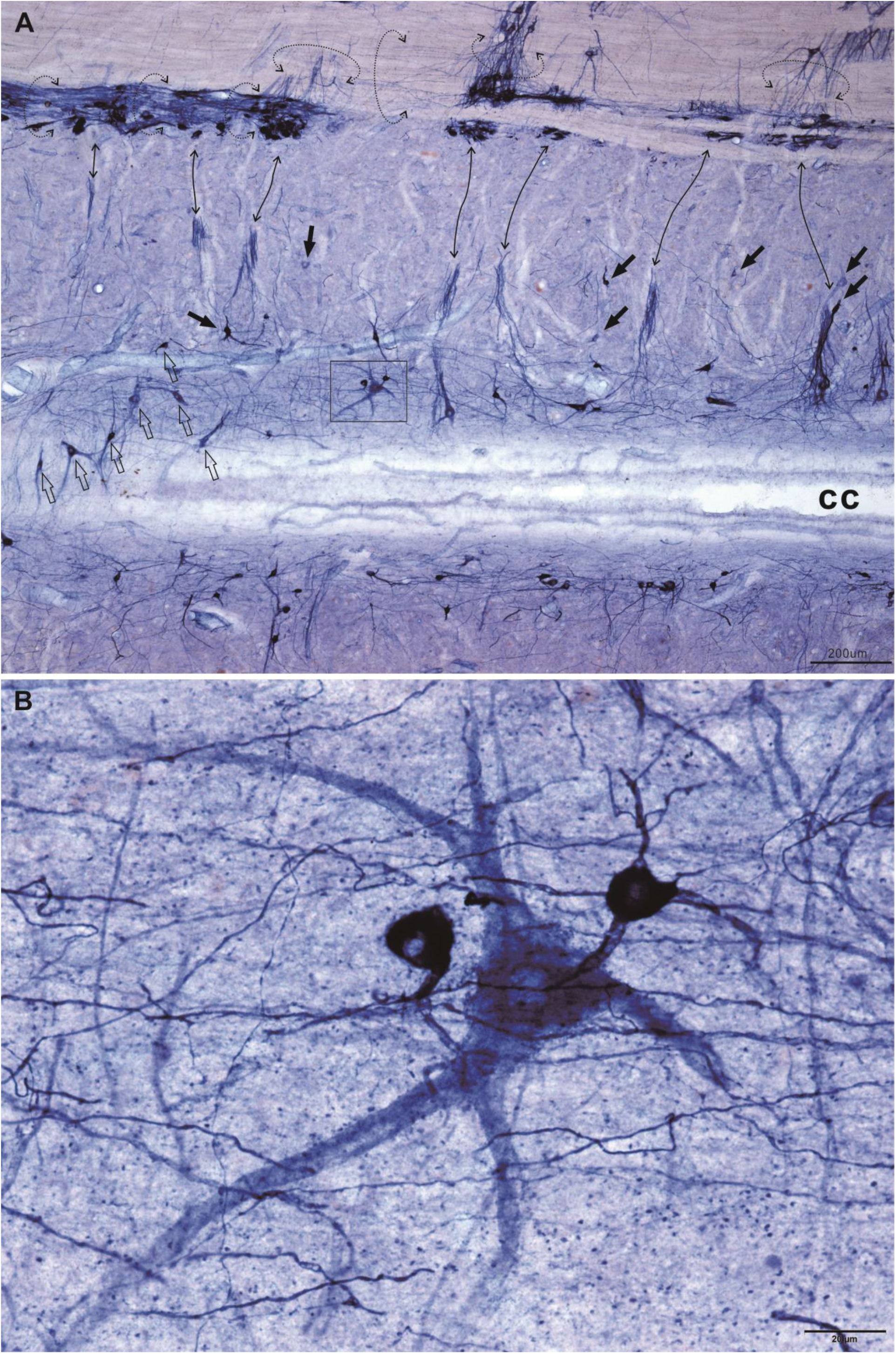
Magnification from Figure 4 to show the IML in the gray matter in the horizontal section(A). Dash circle: fiber bundles. Double arrow: corelated fiber bundle. Arrow: neurons in the intercalated nucleus. Open arrow: neurons in DCN. cc: central canal. B: High power image from A showed two small strong reactive neurons and a large moderate stained neuron. Bar in A = 200μm and bar in B = 20μm.

**Figure 7-1.**
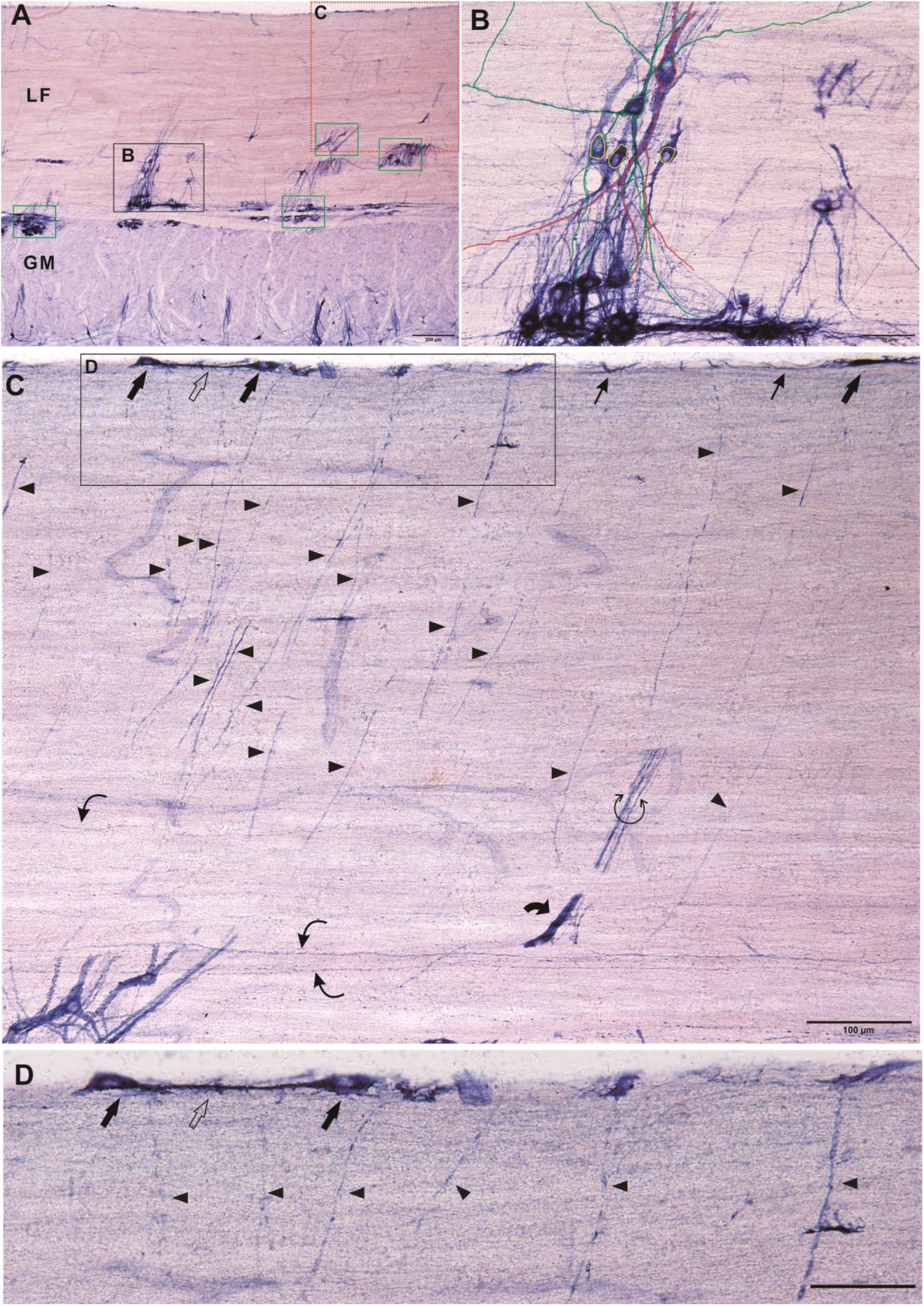
The high-power view of the other side of the lateral funiculus in figure 4. A: The horizontal section of the lateral funiculus (LF) and the gray matter (GM). Green squares will magnify in Figure 7-2. B: Magnification from A showed different profile of the IML neurons and funicular neurons (superimposed color circle): different diameter of the processes and shape of neuronal soma. Red and green lines traced dendrites while yellow one omitted. C: Magnification from A showed pial surface neurons(arrow) and fibers (thin arrow). Arrow: pial neurons. Open arrow and thin arrow: pial fiber. Most of the perpendicular fibers were labeled by arrowhead. Curved arrow: a thick segmental dendrite. Curved thin arrow: longitudinal fibers. Circle arrow: a bundle of fibers. D: Magnification of C. Bar in A 200μm, B and D=50μm and C=100μm.

**Figure 7-2.**
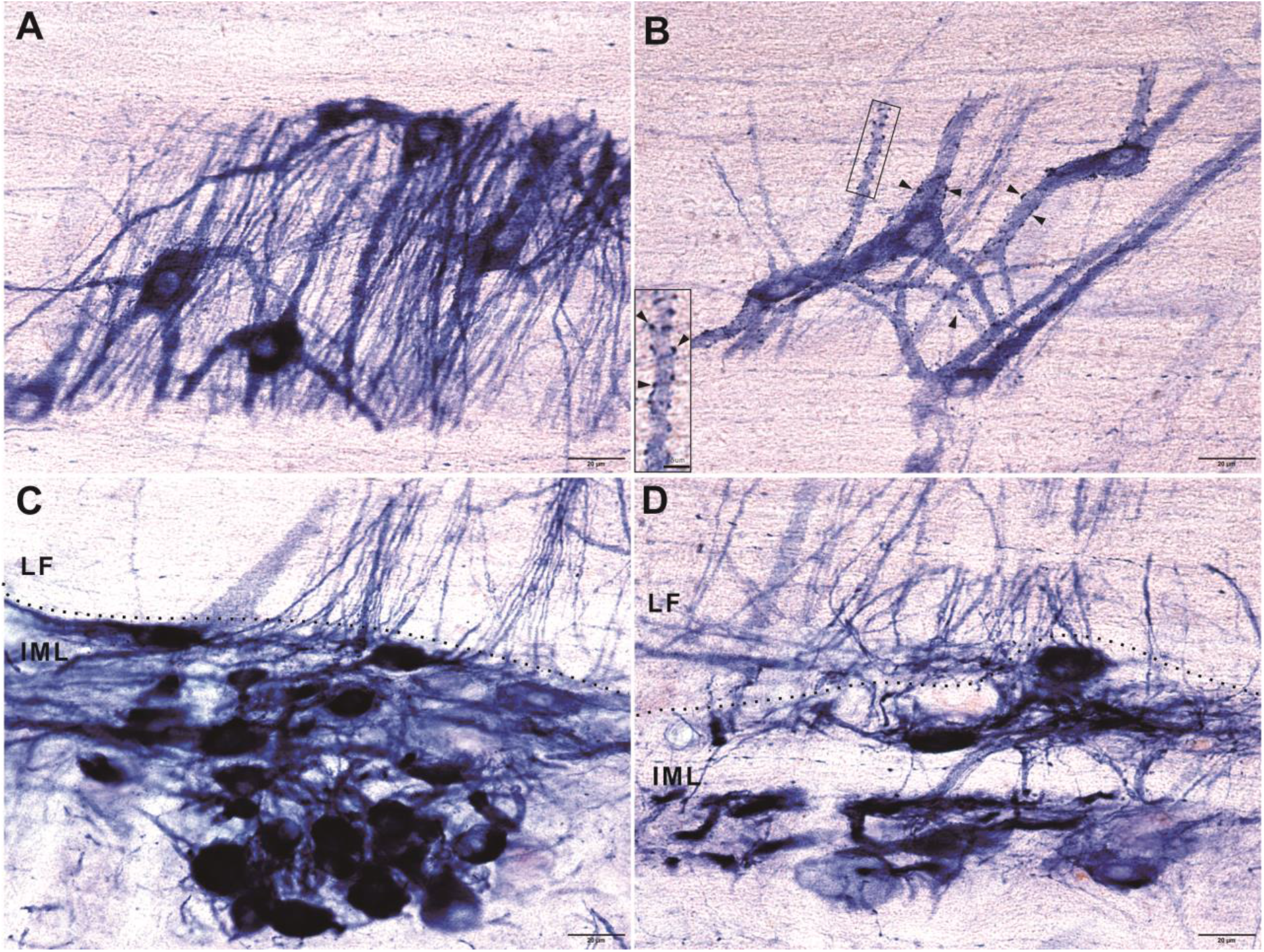
Comparison of IML neurons and funicular neurons. Large soma and multiple dendrites of funicular neurons in A and B. The pericellular particles revealed in the membrane of soma and proximal dendrites (B). The example of pericellular particles indicated both inset and B. C showed the typical IML. Ovary shaped and relatively small neurons in the IML (C and D). The dash line indicated the boundary of the lateral funiculus (LF) and the intermediolateral nucleus (IML). Bar = 20μm and bar of inset = 5μm.

We noticed that the intensity of the NADPH-d staining was different from the medial to lateral white matter. Relative strong staining neurons located in the medial to the gray matter, while the relative light staining neurons located the lateral to the pial surface. It was noted that the relative strong staining showed in the pial surface neurons. This may be related to the external CSF or the divergence and convergence of the projecting dendritic arborization. The pericelluar particle neurons frequently located in the lateral funiculus. There were higher the number of neurons in the medial to the lateral horn of the gray matter.

The funicular neurons and the proximal dendrites were different from those of the neurons in the IML. We made a quantification of the area of soma and the diameter of the proximal dendrites. The area of the neuronal soma in the lateral funiculus was signification larger than that in the IML (p< 0.001). The diameter of the neuron proximal dendrite of funicular neurons was also significantly larger than that of neurons in the IML (p < 0.001) (Figure 8). We also examined other level of horizontal sections. For example, the dorsal horizontal level through the dorsal column still revealed a part of funicular projecting fibers in the white matter. The pial surface neuron was still detected in Figure 9 C and D. The fragment of the funicular projecting fibers occurred in the white matter near the gray matter of the dorsal horn (Figure 9 H).

**Figure 8.**
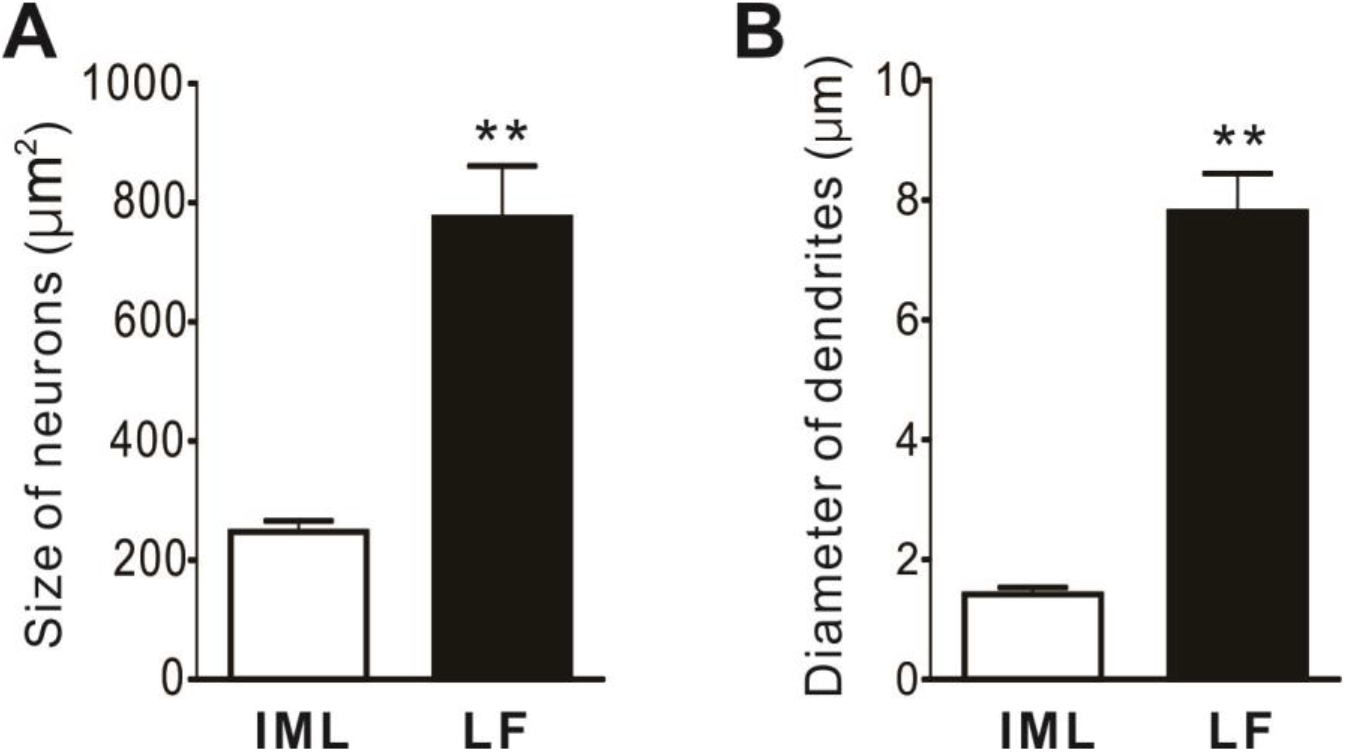
Quantification of the neuronal soma and proximal fibers in the lateral funiculus (LF) and the IML. The area of the neuronal soma in the LF was signification larger than that in the IML. The diameter of the neuron proximal dendrite was also significant between the LF and IML. ** < 0.001.

**Figure 9.**
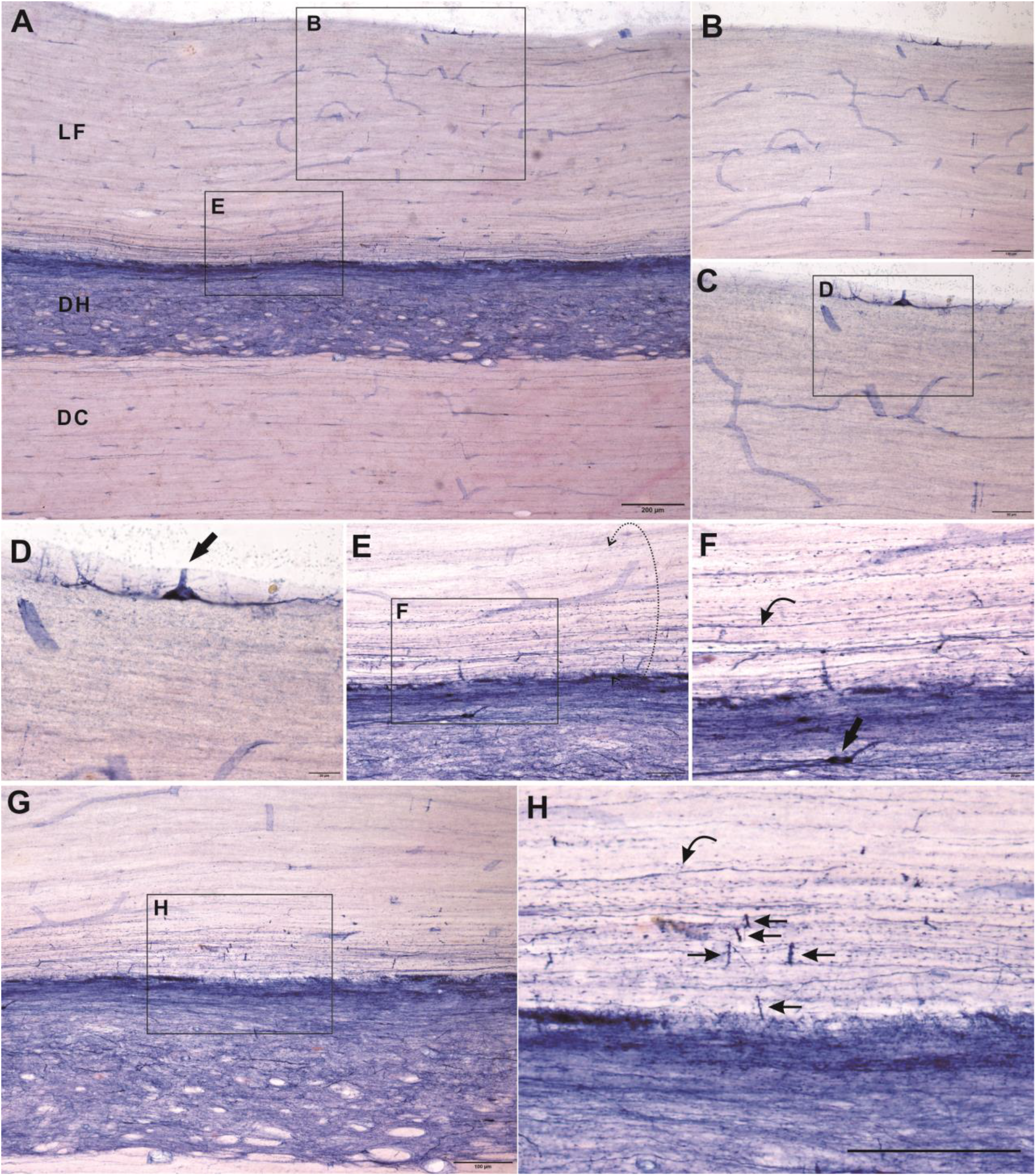
Horizontal section dorsal above IML and lower to the dorsal lateral nucleus as well as through the dorsal column(A). B, C and D showed a neuron located in the pial surface (arrow in D). E showed massive longitudinal fibers indicated by dash circle. An example for fiber indicated by curved thin arrow. Arrow indicated a neuron. G and H showed fragmental fibers (thin arrow) perpendicular to longitudinal fiber (curved thin arrow). Bar in A = 200μm, bar in B, G and H =100μm, bar in C and E= 50μm and bar in D and F = 20μm.

The plexus and cluster of funicular neurons showed partial segmental arrangement along the longitudinal orientation. We figured out a schematic diagram to theoretically illustrate the neuronal configuration of the funicular plexus, subpial plexus and the neurons in the gray matter (Figure 10).

**Figure 10.**
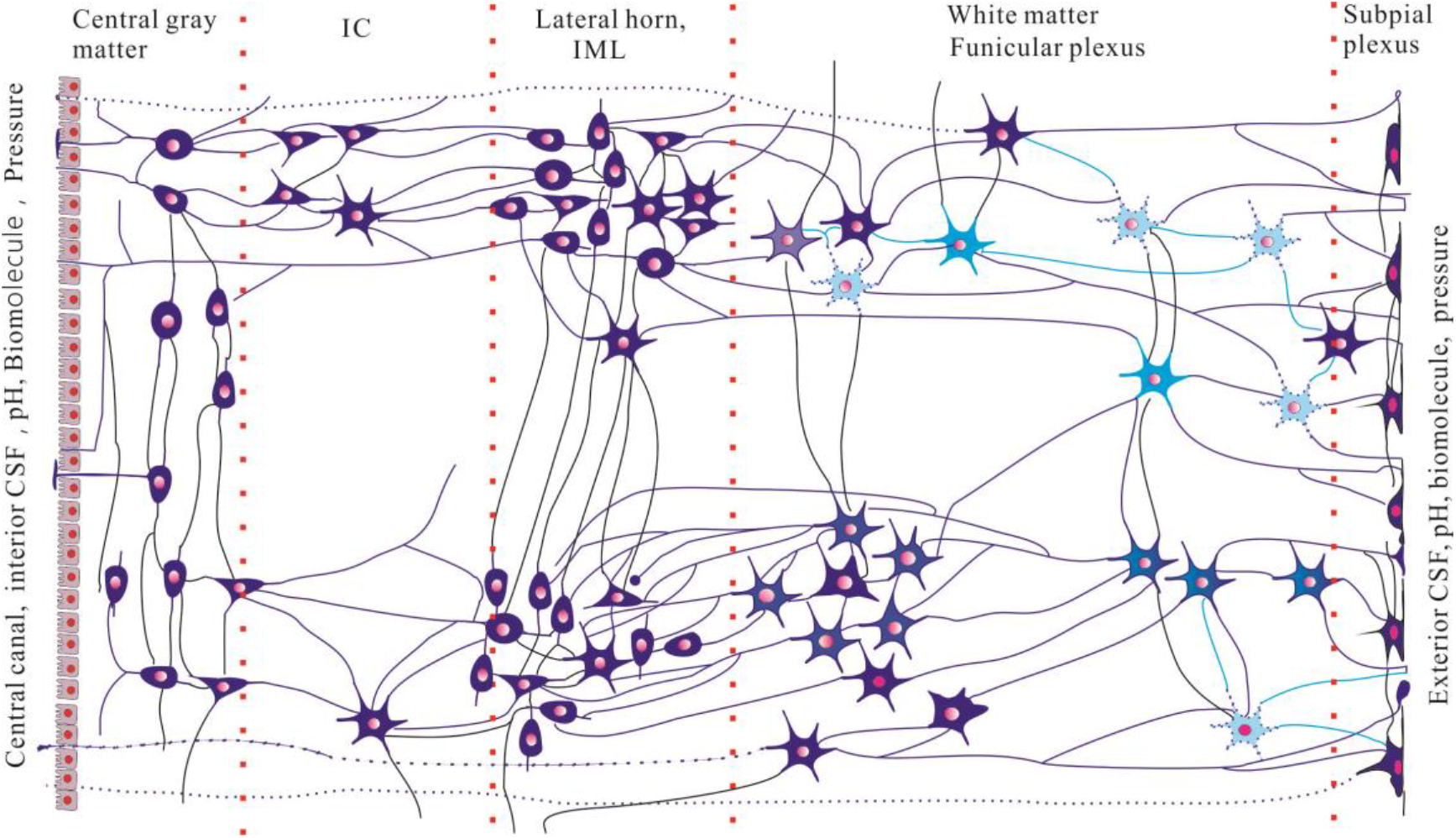
Diagram illustrated the neuronal connectivity in the horizontal arrangement from the central canal to the pial surface. The subpial plexus and funicular plexus reached the external CSF. The pH, bio-molecules and pressure in the CSF may detect by the CSF contacting neuron in the subpial plexus and funicular plexus. IC: nucleus intercalatus spinalis, IML: intermediolateral N.

## Discussion

In this study, we found that cluster of NADPH-d neurons occurred in the mediolateral funiculus. The specialized orientation was typically visualized in the horizontal sections. NADPH-d positivity showed multipolar neurons characteristic of moderately and lightly staining. NADPH-d processes manifested subpial plexus with subpial neurons. It is important for our finding that the funicular neuron and subpial neuron were interconnected and showed reaching CSF in subpial surface. Medial orientation of NADPH-d fibers was also detected cross midline to innervate contralateral side of spinal cord. The subpial plexus were lining surface of subpial region, while the subpial neuronal somas were aligning under pial surface. The plexus and cluster of funicular neurons showed partial segmental arrangement along the longitudinal orientation. For the advantage of detailed visualization of fine terminal, NADPH-d dendritic arborization showed unique configuration completely different Nissl stain[23] or some immunocytochemistry study[30]. The orientation and distributive pattern of the funicular neurons presented a flat-plane organization. The multipolar of the processes clearly showed way of convergence and divergence or single contacting to grey matter with subpial surface. The NADPH-d processes interconnected IML and subpial surface. The surface CSF contacting fibers located caudal of sympathetic segments, which may innervate the pelvic organs[5] and adrenal medulla[22]. With examination of IML in thoracic and lumbosacral spinal cord, we demonstrated that numerous funicular NADPH-d positive neurons and subpial plexus showed in the caudal of sympathetic neurons and relatively less of those were detected in the rostral thoracic segment and sacral spinal cord. Some of IML innervation may not show funicular location[31, 32]. We would like to discuss the funicular innervation and CSF contacting components. It was a location-dependent localization in the white matter.

### Neurons located in the white matter of spinal cord

These NADPH-d neurons and fibers were estimated in the intermediolateral nucleus pars principalis (IMLp) and pars funicularis (IMLf), the nucleus intercalatus, and the central autonomic area[10]. We should term some of IMLf neurons located in lateral funiculus as funicular neuron. Our finding is focused on the funicular neuron. Petras considers IMLf as subpopulation of IML and he also similarly classifies IMLf as preganglionic neuron for its similar profile of somatic motor neuron [23]. The posture-related spinal neurons are considered in the gray matter without examination of lateral funicular neurons in the thoracic segments[33]. In our opinion, profile of each architectonic cell group by NADPH-d histochemistry for funicular neuron is distinguished and different from IMLp. Signaling flow of grey matter was divergence from IMLf to IMLp. Information or inputs from CSF was also convergence from subpial plexus through funicular neurons to IMLp. The cluster of funicular neurons apparently worked as nucleus or regulating center to integrate information from both grey matter and subpial surface. Subpial plexus functioned as receiver(sensor) of CSF. The morphological criteria of subpopulation of those IML were quite different. The soma of large size funicular neurons mostly was a massive junction of giant highways of fibers and the processes of the neurons were also large size compared with IMLp neurons. Funicular neurons were large with thick multipolar wide-spread arborization of processes which set large field of mechanical action. The spatial arrangement of subpial plexus like round brackets located in mediolateral pial surface. These morphological criteria were extraordinary unique feature in our NADPH-d staining. Together with the specialized spatial physical arrangement, we could outline a mechanical and electronic device. Besides autonomic regulation, another function of funicular neurons was supposed to act on movement or balance sensor. Sympathetic neurons in IML works as autonomic output to visceral organs. These neurons are cholinergic neurons identified by choline acetyltransferase (ChAT) immunoreactivity[34–36]. In the white matter of spinal cord, some ChAT positivity are detected in the lateral funiculus[36, 37] and dorsal lateral nucleus[36]. However, a small number of intensely ChAT-positive neurons are detected in the lateral horn of the thoracic spinal cord of the monkey[38]. Ren et al report that a few of Aromatic L-amino acid decarboxylase positive neurons are detected in the lateral white matter in the rat thoracic spinal cord[39]. In some non-mammals, the marginal nucleus locates as a separated nucleus in the white matter[40, 41]. The lateral funiculus regards as origin of sympathetic preganglionic neurons[5, 42, 43]. VIP neurons are also detected in the lateral funiculus [44]. In the upper cervical spinal cord, the white matter neurons project to the intermediolateral cell column[45].

We noted that funicular neurons are investigated with cytoarchitectonics study with a modified Nissl staining in Petras’s experiment [23]. NADPH-d staining in monkey thoracolumbar spinal cord revealed cytoarchitectonics pattern consistent to that of Nissl staining, although the distribution of NADPH-d activity among the species reveals marked different[46].Advantage of NADPH-d histochemistry is NADPH-d positivity displaying fine processes of Golgi-like pattern that shows neuronal cellular connection and fiber projection. While, neurons stained with the Nissl method are individually isolated each other and failed to show fine neuronal processes.

### Subpial region and CSF-contacting neurons in spinal cord

Weakness of our study is that we did not conduct neuronal tracing study. Retrograde tracing can label NADPH-d neurons in both IMLp and IMLf in the upper thoracic cord of rat[47]. And Substance P positive neurons in both IMLp and IMLf are also labeled by retrograde tracing. Interesting for us, subpial plexus and subependymal plexus are labeled by cholera subunit B conjugated horseradish peroxidase (CB-HRP)[48]. It should be important that the widespread CSF contacting or subpial reaching fiber take integrating information with such anatomy configuration. However, it is hardly concluded that subpial plexus is directly correlated with NADPH-d subpial plexus. We did not note similar subpial plexus in the sacral spinal cord by HRP neuronal tracing[48, 49]. Our unpublished data also showed that CB-HRP could label subpial terminals in rat and chicken sacral spinal cord for pelvic organs. It may be different of HRP tracing for CB-HRP tracing in different animal study[49]. Substance P and NADPH-d positivity in IML are changed in the hypertensive animals[47, 48]. LaMotte reports that vasoactive intestinal peptide (VIP) contacting neurons occur the central canal and also along with subpial surface in the thoracic and sacral spinal cord of monkey and cat[50]. Enkephalin positive fibers are also detected in the lateral funiculus of rat [51]. Recently, we reported that VIP neurodegenerative megaloneurites are detected in the sacral spinal cord in the aged dog[15]. It demonstrates that both VIP and NADPH-d CSF contacting structures exist in the spinal cord. In the monkey and cat, Lamotte reports that VIP CSF contacting positivity is rich in spinal central canal[52]. Djenoune introduces that Kolmer and Agduhr have identified and described spinal CSF-contacting neurons in over 200 species[28]. Vigh reports that there are terminal enlargements at the ventrolateral surface of the spinal cord studied by Golgi method based on study of a total of 136 submammalian vertebrates[53]. There is nerve terminal area formed by the axons of the CSF contacting neurons in ventrolateral spinal surface[53–55]. Different to the submammalian vertebrates, our data showed chemical and function morphological property in the subpial surface of the spinal cord by the means of NADPH-d enzyme histology. We thought that NADPH-d funicular neurons played more important role in the non-human primate for the specialized CSF pathway. NADPH-d may also regulate cerebral blood flow through CSF in ventricle[56]. The NADPH-d funicular neurons of subpial plexus may not directly contact with central canal instead of IMLp and also can make circuit with neighbor neurons. In the lamprey spinal cord, two types of CSF contacting neurons can be labeled from the subpial region[57]. Comparison investigation from fish to primate, Djenoune et al think that the CSF contacting neurons locate the central canal area after examined the cervical, thoracic and lumber spinal cord of the monkey [58]. It is the first time for us to demonstrate CSF contacting neurons and fiber through subpial plexus in the thoracolumbar spinal cord of the non-human primate.

CSF-contacting neurons regulate motor control[59, 60]. Böhm recently reports that CSF-contacting neurons form an intraspinal mechanosensory organ that is relevant to locomotion because it [61]detects spinal bending[62]. Hubbard et al report that intraspinal sensory neurons modulate motor circuits of locomotion[63]. Most of the experiments are completed in fish[64]. Relatively, the spinal cords of birds have a glycogen body not seen in mammals[65].The glycogen body may work as a sense organ of equilibrium [66]. We try to demonstrate the structure with NADPH-d histology[16]. There are only a few of NADPH-d fiber around in the structure. Harkema et al report that epidural stimulation of the lumbosacral spinal cord facilitates patient of motor complete paraplegia with assisted stepping[67].

CSF-contacting neurons in spinal cord are also act as pH sensors[68, 69]. We postulated that sympathetic neurons need information of CSF to maintain homeostasis of brain circumstance. The caudal portion of CSF may be important for sympathetic regulation. Cluster of funicular neurons may play important role of feedback information for CSF circulation. Inner humoral inputs from CSF for sympathetic preganglionic neurons was a critical regulating loop for CSF system, although sympathetic preganglionic neurons are output with distinct spinal origin[5].

Vera raises an interesting question:“the intermediolateral nucleus: an’ open’ or’ closed’ nucleus’?”[32]. From this point of view, we made up that the NADPH-d funicular neurons may be considered “open” to CSF. In order to understand the funicular neurons and subpial plexus for CSF contacting, subcommissural organ and CSF circulation were briefly reviewed. SCO-Reissner’s fiber complex is one of the factors involved in the CSF circulation [70]. CSF for the delivery of various factors throughout the central nervous system[71]. We may need more devices to maintenance autonomic control of CSF in caudal anatomical position. Differently, most of CSF contacting neurons or SCO and other periventricular structures are located in the ventricular system and central canal[72–76]. Our finding revealed a structure out of ventricular structures. Funicular neurons and subpial plexus could make more precise regulation of CSF maintenance. To a certain extent, related structure for maintenance of CSF may be relevant to neurogenesis[77]. We may also postulate that degeneration of NADPH-d funicular neurons could cause sensory-somatic dysfunction with aging, if it functioned as mechanical sensor. We do find that NADPH-d dystrophic neurites occur around the central canal in aged dog[15]. Next in our future experiments, we should identify Pkd2l1 expression in the NADPH-d funicular neurons for its maintenance of spine curvature[60]. In aged dog, Reissner’s fiber, a “central body” or a “bud” [27]could be NADPH-d positive reaction[15]. In relative lower vertebra, such as fish, there is a specialized structure, urophysis related to neurosecretory cells[78]. In higher order species of vertebrae, the CSF contacting fiber also is found to reach the external CSF [72]. For terminology, some investigators also use liquor-contacting neuron for the CSF contacting neuron[79]. This clue reminded us that the present circuits in white matter in thoracolumbar spinal cord may also serve as neuroendocrine apparatus in the non-human primate, especially relevant to subpial plexus.

**Terminology:** pial surface CSF-contacting texture, subpial and funicular plexus or Tan plexus

It is noted that there are several neurons located in the dorsal column white matter of the spinal cord in several mammals, including primates and birds. These neurons play potential function and can response to bioactivity change in CSF [80]. We should design similar experiment to test funicular neurons in future. Compared with our finding, the neurons in dorsal column are relatively and individually separated and scattered distribution. A crucial configuration for the lateral funicular neurons showed not only a pattern of clusterization and plexus, but also horizontal section through the central canal, lateral horn and lateral funiculus in our data. But we should distinguish the lateral funicular neurons from the nucleus in the dorsolateral funiculus of the spinal cord of the rat[81]. We are more familiar with the lateral horn, in which the number of neurons is analysis for study of autonomic system[3, 82].

“Plexus” was chosen as term to name our preliminary finding for the anatomical profile of the cluster of the funicular neurons is similar myenteric plexuses: myenteric plexus and submucosal plexus. It is important of regulation also called autonomic plexus or enteric plexus. Other terms for myenteric plexus and submucosal plexus are Auerbach’s plexus [83] [84]and Meissner’s (plexus) research[84] which name the terms after researchers. We would like to discuss several cases for using “plexus”. “The motoneuron plexus” and “longitudinal plexus” are used in anatomical organization of the spinal cord in cats[85]. In this paper, the motoneuron plexus is preference for function and the longitudinal plexus is preference for anatomical orientation. Barber demonstrates that some ChAT neurons terms partition neurons distribute in “longitudinal plexus”[37]. “Neuroplexus”[86–88] are frequently used gastroenteric study or eye and vision science. Another reason for using “plexus” is that choroid plexus is a common anatomical term[89]. “The lateral plexus” is used in the retrograde labeling of the CSF contacting neurons in the lamprey spinal cord[57]. The location of the lateral plexus is similar to our subpial plexus. Definitely, we considered “ganglia” as term to name our finding. Readers should prevent funicular plexus and subpial plexus from the nucleus in the dorsolateral funiculus of the spinal cord of the rat[81].

Petras reports that IML neurons consistent of IMLp and IMLf which they abbreviated ILp and ILf[23]. They demonstrate that many neurons of IMLp and IMLf are preganglionic neurons. The size of the neurons is as large as somatic motor neurons. But we found that the size and shape of IMLf were not exactly the same as that of IMLp. The striking feature of funicular neuron is flat-oriented soma interconnected cluster like ganglia. The individual soma and the cluster of neurons were orientally vertical to the IML. We thought that IML is anatomical term of nucleus in the gray matter. IMLf involves neurons in the whiter matter. The “lateral” for IML (intermediolateral) is the neurons located lateral of the gray matter. The distribution of peptide-like immunoreactivity in IMLf is quite different[10, 90]. We focused on funicular neurons which can group as ganglionic pattern and connect with processes as plexus. Straightforward, we may suggest to name funicular plexus and subpial plexus after researcher’s last name to some extent. In order to prevent the confusion, Tan plexus was considered for funicular plexus and subpial plexus in the lateral funiculus of the spinal cord of the monkey. The dendritic arborization of IML neurons labeled by neuronal tracing is radial to white matter[91]. The neural circuit of kidney shows the innervation of neurons located in the both IMLp (IML) and IMLf (lateral funiculus) in thoracic spinal cord [92]. However, the double staining neurons of NOS and tracing labeling majorly locate in the IMLp. The central sympathetic circuits regulating viscera and the spinal sympathetic reflexes are mediated polysynaptically[93, 94]. The preganglionic neurons in IMLp connect with interneurons[95], however the funicular neurons may have a different way to make a circuit with interneuron[96]. But some of the funicular neurons clearly contacted with NADPH-d buttons or punctas in our present finding. The NADPH-d funicular neuron formed triangle radiated dendritic arborization majorly directed to IML and subpial plexus. Not all the researchers identify the IMLf as anatomical term to present their research data[20, 97]. In our data, the pericellular punctate NADPH-d particles surrounding neurons were strikingly different from the neurons located in the lateral horn[98]. This is similar to our recent claimed NADPH-d pericyte in the medulla oblongata in the pigeon. NADPH-d staining is used in “the Rat Brain in Stereotaxic Coordinates” and “Chemoarchitectonic Atlas of the Rat Brain” as a standardized staining method for cytoarchitecture[99–101]. The IML in the thoracic cord of the rodent may be not consistent with that of the non-human primate[102]. We name the NADPH-d funicular neurons as funicular plexus or Tan plexus based on the cytoarchitecture by NADPH-d staining which are widely used for myenteric plexus[103, 104].

The angular distribution of neuronal texture was supposed a directional dependence of motion sensation. Finally, we had a hypothesis that the funicular neurons may detect and react to the motion of spinal cord when body truck in flexion/extension, lateral bending, or axial rotation based on the cytoarchitectonics, besides the detection of the pressure and biochemical environmental change of the CSF. We make an illustration to demonstrate the hypotheses (Figure 11).

**Figure 11.**
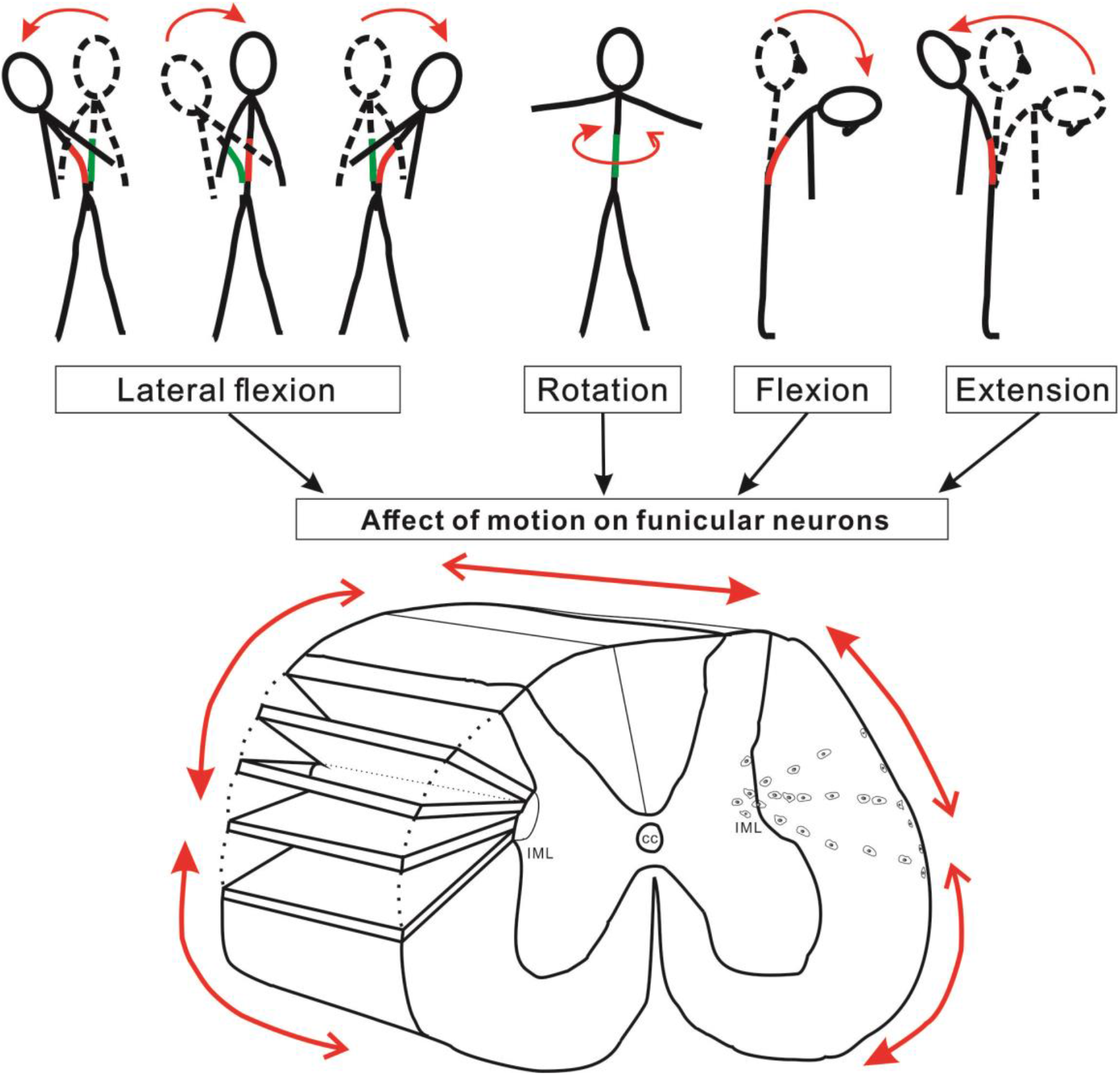
Schematic diagram of the funicular neuron mechanosensory modulation in the thoracolumber spinal cord. Arrows indicated the direction related motion pattern and posture.

## Summary

Morphological feature and location of neuron should be a gold standard to make some inferences according to our observations. Nissl staining shows isolated neuronal cells. With advantage of NADPH-d, the detailed neuronal dendrites were well visualized. We first to reveal neuronal texture by NADPH-d. The funicular neurons with fiber distribution and anastomose were termed academically as funicular plexus and specialized localization for subpial structure we termed subpial plexus. We did not find the NADPH-d funicular neurons in the sacral spinal cord and rostral of thoracic spinal cord was not found obvious NADPH-d funicular neurons and subpial plexus. We thought that the location of the funicular neurons and subpial plexus set a suitable position for detecting active bending of the spinal cord. It may help several following actions: detecting body position, circulation and drainage of CSF and sensation of molecule change in CSF. Djenoune demonstrates that CSF-contacting neurons is a sensory interface linking the CSF to motor circuits in vertebrates[26]. We found that predominantly interconnected NADPH-d funicular neurons contacting sympathetic neurons. Summarily, the CSF contacting NADPH-d funicular neurons and subpial plexus could be a structure to regulation of sympathetic output for somatic motion or regulating visceral organs and ensure CSF homeostasis.

## Acknowledgments

This work was supported by grants from National Natural Science Foundation of China (81471286), Liaoning Training Programs of scientific research and career development for Undergraduates, 201410160007 and Research Start-Up Grant for New Science Faculty of Jinzhou Medical University.

